# Cryo-EM and single molecule visualization unravel the role of human condensin II activation by M18BP1 in driving DNA compaction

**DOI:** 10.64898/2026.02.02.703226

**Authors:** Alessandro Borsellini, Sebastian Chamera, Valentina Cecatiello, Joanna Andrecka, Alessandro Vannini

## Abstract

DNA compaction by human condensin II is crucial for the correct organization of mitotic chromosomes. Mitotic activation of condensin II is a multi-layered process, controlled mainly by the opposing activity of the MCPH1 and M18BP1 proteins, acting as an interphase inhibitor and a mitotic activator, respectively. Mitotic phosphorylation favors M18BP1 binding to condensin II, resulting in effective and timely chromosome compaction. However, the molecular mechanisms underlying condensin II activation in driving DNA compaction remain uncharacterized. Combining cryo-electron microscopy with single molecule imaging we unravel the mechanism of condensin II activation by M18BP1 and elucidate their role in DNA compaction. The specific interaction with phosphorylated M18BP1 induces conformational changes in the condensin II complex, relieving an autoinhibitory conformation and thus allowing stable DNA binding and DNA compaction. Beyond the ATP-driven activity of isolated condensin II complexes on DNA, we observe the emergence of multimeric species that withstand the forces applied by optical tweezers, highlighting the contribution of condensin II protein–protein interactions to the mechanical reinforcement of mitotic chromosomes.

## Main Text

At the heart of eukaryotic cell division lies the dramatic reorganization of the chromatin into compact, rod-shaped chromosomes that can be faithfully divided between the daughter cells. An important step to achieve such reorganization is the volumetric compaction of the chromatin favored by histone post translational modifications like phosphorylation and de-acetylation (*1, 2*). On top of the volumetric compaction, a series of mechanical activities carried out by condensin I and II, in concert with Topoisomerase IIα and the chromokinesin KIF4A are fundamental to organize the sister chromatid axes contributing to stiffness and compaction (*3-7*). Condensin I and II, both members of the Structural Maintenance of Chromosomes (SMC) family, shape the higher-order architecture of chromosomes by forming chromatin loops, however increasing evidence indicates that additional mechanisms, might contribute to mitotic chromosome assembly (*8-12*). While condensin I and II share the core SMC2–SMC4 heterodimer, they differ in their sets of HEAT-repeat subunits - CAP-D2, CAP-G, and CAP-H in condensin I, and CAP-D3, CAP-G2, and CAP-H2 in condensin II - which provide distinct regulatory and functional properties. Condensin II, which resides constitutively in the nucleus, initiates chromatin loop formation in prophase, setting the stage for the nested loop architecture further refined by condensin I upon nuclear envelope breakdown (*13-16*).

Given its nuclear presence throughout the cell cycle, condensin II is subject to stringent regulation to ensure that chromosome condensation begins precisely at mitotic entry. Two major mechanisms have been identified thus far: first, condensin II is activated at mitosis through the interaction with the phosphorylated form of the M18BP1 protein that outcompetes the inhibitory protein MCPH1 (*17-19*). MCPH1 prevents premature condensin II activity during interphase. Secondly, mitotic phosphorylation of the condensin II CAP-D3 tail (residues 1297-1498) relieves an auto-inhibitory interaction with the CAP-G2 subunit, enabling full activation of condensin II (*20*).

While the role of M18BP1 as an essential factor that triggers condensin II activity at the onset of mitosis has recently been established (*18*), the molecular mechanisms underlying the M18BP1-dependent activation of condensin II to initiate DNA compaction are still elusive. By combining single-molecule optical tweezers and cryo-electron microscopy, we elucidate the mechanism of condensin II activation by M18BP1 at the molecular level, unraveling distinct conformational states that underlie condensin II function. Binding of phosphorylated M18BP1 to condensin II stimulates the formation of stably DNA bound complexes capable of DNA compaction. Phosphorylated M18BP1 binding induces extensive structural rearrangements of the condensin II complex, specifically relieving an autoinhibitory interface to allow productive DNA clamping and compaction.

## Results

### Phosphorylated M18BP1 promotes stable condensin II – DNA interaction

To investigate whether phosphorylated M18BP1_873-1132_ (M18BP1^P^) modulates the DNA binding properties of condensin II, we employed a dual-trap optical tweezers system with confocal fluorescence detection, allowing direct visualization and quantification of individual fluorescently labelled proteins interacting with DNA (Fig. 1A) (*21-23*). Pre-incubation of condensin II with M18BP1^P^ increased both the number of binding events as well as the dwelling time of condensin II on the DNA molecule (Fig. 1B-D). The observed enhanced DNA-binding activity is the direct consequence of the formation of stable condensin II-M18BP1^P^ complexes, since doubly labelled condensin II-M18BP1^P^ complexes colocalize on DNA and the M18BP1^P^ protein does not bind to DNA in isolation (Fig. S1). Quantitative analysis of kymographs reveals M18BP1^P^ binding nearly doubled the mean residence time of condensin II on DNA (5.3 s ± 0.5 s to 9.4 s ± 0.6 s) (Fig. 1B,C). Mean square displacement (MSD) analysis reveals that condensin II in presence of ATP exhibits dynamic movement suggestive of one-dimensional diffusion and transient engagement with DNA (Fig. 1B,D). In contrast, the interaction with M18BP1^P^ appears to limit the mobility of condensin II. The diffusion coefficient of condensin II decreases almost 10-fold in the presence of M18BP1^P^ (0.11 ± 0.01 μm^2^/s to 0.01 ± 0.003 μm^2^/s), indicating a more stable condensin II-DNA interaction upon M18BP1^P^ binding (Fig. 1B,D). As a control, a M18BP1 mutant lacking the phosphorylation sites required for condensin II binding (M18BP1^7A^) (*18*) did not alter the mode of condensin II binding to the DNA, which remained diffusive and transient with a diffusion coefficient similar to the one observed for condensin II in isolation (0.11 ± 0.01 and 0.09 ± 0.01 μm^2^/s, respectively) (Fig. 1E).

**Fig. 1.**
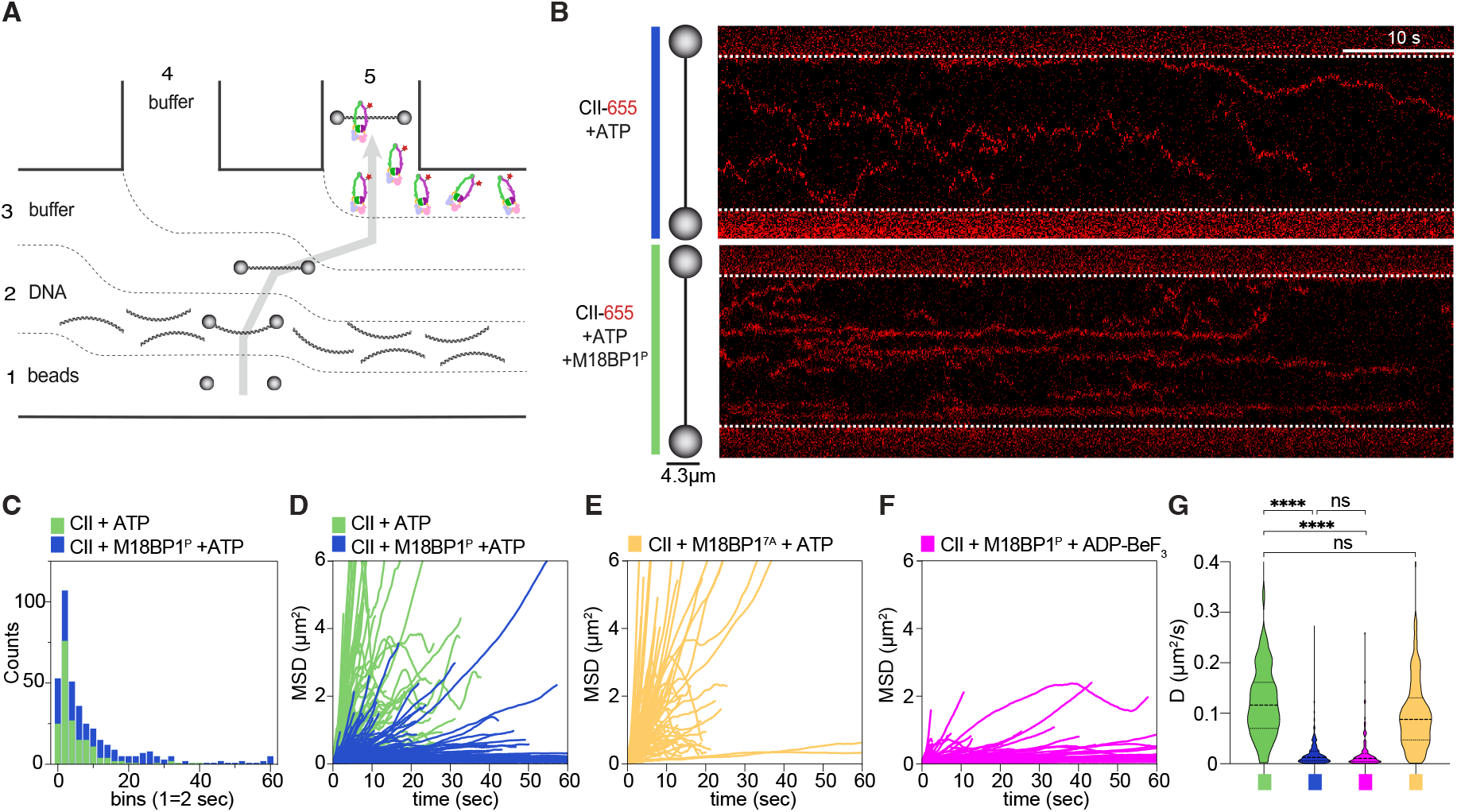
Condensin II loading on DNA is enhanced by M18BP1. (**A**) Schematic representation and channel composition of the microfluidic device used in the optical tweezers experiments. Channels 1 and 2 were used to trap streptavidin-coated polystyrene beads (4.1 µm diameter) and single biotinylated λ-DNA molecules (48.5 kb), respectively; Channel 3 and 4 contained experimental buffer; Channel 5 was used for protein loading and imaging of binding events. Gray arrow represents the path of the beads throughout the experiment. (**B**) Representative kymographs imaged in the protein channel. Red traces indicate the position over time of LD655-labelled condensin II molecules. In the top panel, 0.5 nM LD655-labelled condensin II was incubated with λ-DNA in presence of 3 mM ATP. In the bottom panel, 0.5 nM LD655-labelled condensin II mixed with 20-fold excesses of M18BP1^P^ was incubated with the λ-DNA in presence of 3 mM ATP. Kymographs were recorded for 80 seconds. (**C**) Histogram from 8 different kymographs of condensin II DNA binding events in presence (blue) and absence (green) of M18BP1^P^. (**D**) Mean squared displacement (MSD) plots of condensin II trajectories along the DNA in presence (blue) and absence (green) of M18BP1^P^. (**E**,**F**) Mean squared displacement (MSD) plots of condensin II trajectories along the DNA in presence of M18BP1^7A^ and ATP (E, yellow) or M18BP1^P^ and ADP-BeF_3_ (F, magenta). (**G**) Violin plot representing the distribution of Diffusion Coefficients extracted from the MSD data in the four different conditions shown in panel D-F.

In order to determine whether ATP hydrolysis by condensin II is required for the observed stable DNA binding mode, condensin II was incubated with M18BP1^P^ and ADP beryllium fluoride (ADP-BeF_3_), which locks the ATPase domains in a pre-hydrolytic state (Fig. 1F). Stable and non-diffusive DNA binding by condensin II was observed with a diffusion coefficient similar to the one observed in presence of M18BP1^P^ and ATP (0.009 ± 0.004 μm^2^/s) (Fig. 1F). Taken together, these experiments demonstrate that M18BP1^P^ profoundly alter condensin II-DNA interactions, triggering the formation of abundant stable non diffusing condensin II - M18BP1^P^ - DNA complexes, which do not require ATP hydrolysis for their association with DNA (Fig 1G).

### Stable condensin II – M18BP1^P^ complexes induce DNA compaction

Next, we asked whether the stable M18BP1^P^-condensin II complexes could induce DNA compaction, and performed relax-stretch assays (Fig. 2A, Fig. S2A), as previously described for meiotic protein SYCP3 and yeast cohesin (*24, 25*). In presence of ATP, stable condensin II - M18BP1^P^ complexes consistently generated force-distance profiles indicative of DNA compaction, whereby the measured force between the beads increased at lower distances with a distinctive sawtooth shape (Fig. 2B-D). Interestingly, the fluorescent traces that are initially spatially separated along the DNA consistently converge into a single, more intense fluorescent trace (Fig. 2C, Fig. S2B-D). This phenomenon suggests that condensin II molecules, in the presence of ATP, M18BP1^P^ and when bound to relaxed DNA molecules, coalesce together to form individual, more stable protein-DNA clusters that withstand high forces. Imaging the same experiments in wide field modality reveals the fast formation of individual protein-DNA clusters that cannot be resolved by applying high forces (Fig. 2E). In absence of M18BP1^P^, no DNA compaction activity was detected, and the resulting force-distance curves did not deviate from the ones obtained in presence of the λ-DNA molecule in isolation (Fig. S3A). Importantly, the ability to form multimeric protein–DNA clusters appear to be conserved among human condensins. In the same experimental setting, condensin I compacted DNA and formed stable protein–DNA clusters similarly to what observed for the condensin II – M18BP1^P^ complexes (Fig. S3B), in agreement with recent data showing condensin I mediated “DNA lumps” formation in single molecule assays (*26*). Thus, while M18BP1^P^ binding is required to restructure condensin II into a DNA compaction-competent complex, condensin I is able to constitutively compact DNA in absence of auxiliary factors.

**Fig. 2.**
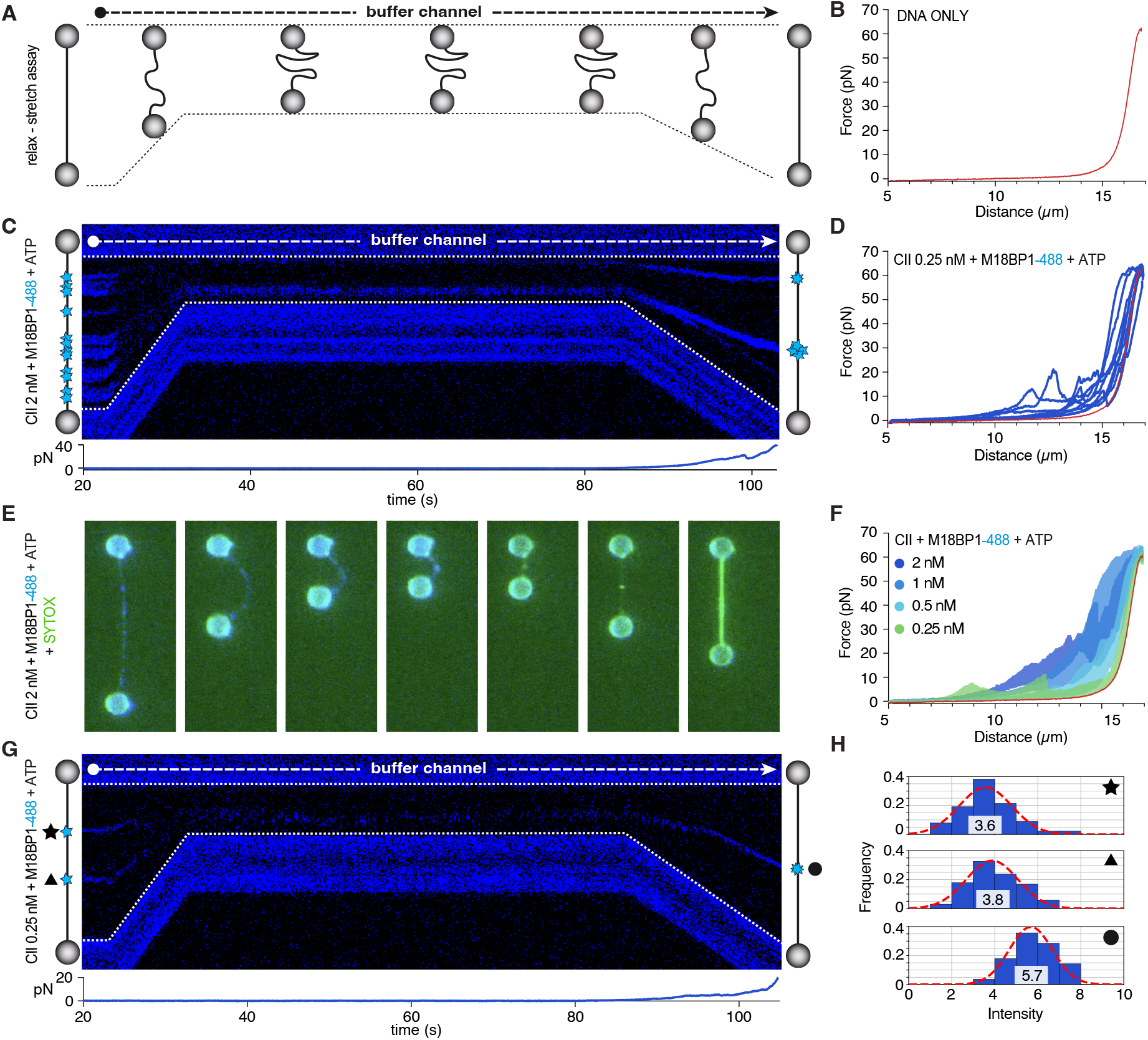
Stable condensin II – M18BP1^P^ complexes compact the DNA. (**A**) Schematic of the relax-stretch assay. After observing DNA binding events in the protein channel, the tethered λ-DNA is transferred to the buffer channel, ensuring no additional DNA-binding events (Fig. S2A). In the buffer channel, the distance between the beads is reduced from 12 µm to 5 µm allowing relaxation of the λ-DNA molecule. At 5 µm distance, the DNA and bound proteins are incubated for one minute in presence of 3 mM ATP, after which the DNA is stretched to 17 µm while monitoring force-distance curves. (**B**) Force-distance curve of λ-DNA molecule in absence of proteins. (**C**) Kymograph of the relax-stretch assay performed in presence of 1 nM condensin II, 10 nM M18BP1^P^ (AF-488) and 3 mM ATP. Below the kymograph, a plot of the respective force recorded during the experiment is shown. (**D**) Force-distance curves of λ-DNA molecule from 8 replicates of experiment shown in panel C. (**E**) Snap shots of relax-stretch experiment performed in same conditions as experiment in panel C but imaged using wide field microscopy. Light blue signal comes from M18BP1^P^ proteins labelled with AlexaFluor488, while green signal comes from SYTOX dye that intercalates the DNA molecule. (**F**) Force-distance curves of titration experiments. Each condition is the average of 5 replicates. Dark blue represents highest concentration (2 nM) while light green represents lowest concentration (0.25 nM). (**G**) Representative kymograph of relax-stretch assay performed with condensin II 0.25 nM and M18BP1^P^ 10 nM and ATP 3 mM. Below the kymograph, a plot of the respective force recorded during the experiment is shown. (**H**) Quantification of fluorescence intensity of traces from kymograph shown in panel G, reporting mean values and Gaussian fits. Black symbols indicate the trace of the kymograph from which the histogram were extracted.

### Two modes of DNA compaction

The stretch-relax assays carried out in presence of condensin II and M18BP1^P^ displayed consistent shortening of the λ-DNA contour length (Fig. 2C-F), representative of persistent compaction events that are resistant to the applied force. In addition, smaller sawtooth-shaped peaks could be observed at shorter distances which might indicate rupture of DNA loops (*24, 25*). To further characterize the DNA compaction behavior observed in the stretch-relax assay in presence of stable condensin II-M18BP1^P^ complexes, we performed a serial dilution experiment of condensin II, (ranging from 0.25 nM to 2 nM), in presence of a fixed concentration of M18BP1^P^ (10 nM, Fig. 2F). Even at the lowest concentration probed, whereby individual condensin II-M18BP1^P^ complexes are spatially separated by several kilobases, the force distance profiles display appreciable DNA compaction features (Fig. 2F-H, Fig. S3C,D). As expected, the degree of DNA compaction increases proportionally with the number of stable complexes bound to the DNA at higher concentrations (Fig. 2F, Fig. S3C,D). To probe whether DNA compaction requires condensin II ATPase activity, a previously characterized ATPase-deficient mutant (condensin II^QL^) (*27*), was employed. Surprisingly, while at low protein concentrations condensin II^QL^-M18BP1^P^ complexes were unable to drive DNA compaction, an appreciable degree of DNA compaction could be detected at higher protein concentrations even in absence of ATP hydrolysis (Fig. S4). A similar behavior was observed for wilt-type condensin II - M18BP1^P^ in presence of ADP-BeF_3_ (Fig. S4). Taken together, these results suggest that two distinct mechanisms contribute to the DNA compaction observed in the stretch - relax assay: (1) an ATP hydrolysis-dependent mechanism, that enables individual condensin II complexes to compact the DNA, associated with reversible rupture events and (2) an ATP hydrolysis-independent bridging mechanism between condensin II– M18BP1^P^ complexes that likely compacts the DNA when the bound proteins are in close proximity, associated with persistent compaction events.

### Cryo-EM structure of the condensin II – M18BP1^P^ clamped DNA complex

Single molecule assays demonstrated that the binding of M18BP1^P^ strongly activates condensin II binding to DNA. To understand how M18BP1^P^ activates condensin II, we used cryo-electron microscopy. The condensin II – M18BP1^P^ complex was incubated with 200-bp double-stranded DNA (dsDNA) and mixed with the non-hydrolysable ATP analogue ADP-Beryllium Fluoride (ADP-BeF_3_) before crosslinking with Sulfo-SDA. By keeping the concentration of the sample relatively low before vitrification it was possible to reduce the tendency to form aggregates, and individual particles isolated from the micrographs were refined at 6 Å average resolution (Fig. 3A, Fig. S5A-G).

**Fig. 3.**
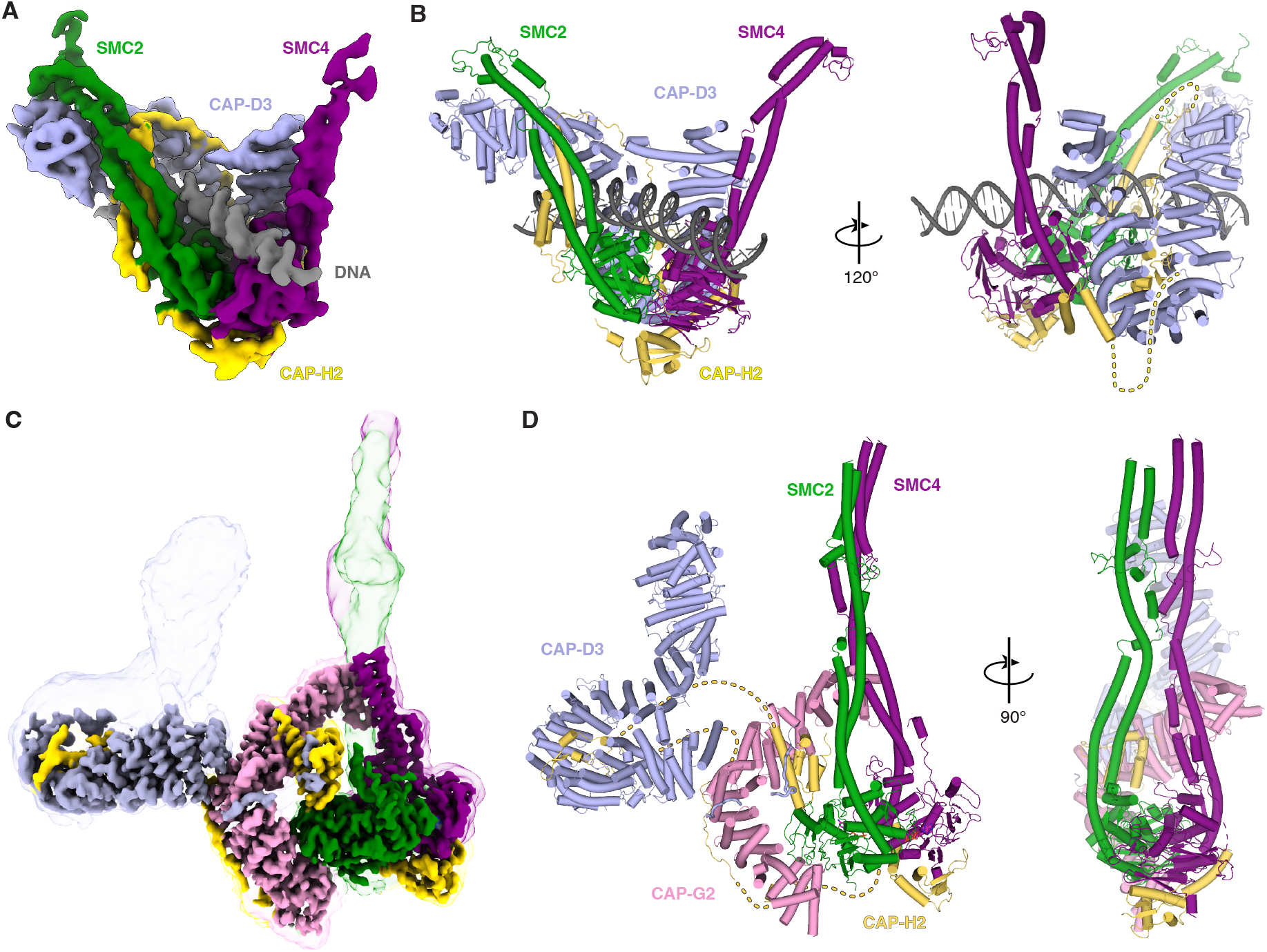
Cryo-EM structures of condensin II complexes. (**A**) Cryo-EM map of condensin II-M18BP1^P^ complex bound to clamped DNA. SMC2 is colored in green, SMC4 in purple, CAP-D3 in pale blue, CAP-H2 in yellow, and the DNA in grey. (**B**) Model of condensin II bound to clamped DNA, colored as in panel A. Dotted yellow lines represents the approximate position of the CAP-H2 disordered regions. (**C**) Cryo-EM map of condensin II bound to two ADP-BeF_3_ molecules. SMC2 is colored in green, SMC4 in purple, CAP-D3 in pale blue, CAP-G2 in pink and CAP-H2 in yellow. (**D**) Model of condensin II bound to two ADP-BeF_3_ molecules colored as in panel C. Dotted yellow lines represents the approximate position of the CAP-H2 disordered regions.

The resulting structure displays typical features of DNA-bound SMC complexes, with the SMC2 and SMC4 ATPase domains, CAP-D3 and CAP-H2 that form a “clamped head module” (*28-30*). The SMC domains are connected through the ATPase heads and the coiled coils are open in divergent directions (Fig. 3B). The DNA lies on top of the ATPase heads and is held in place by the CAP-D3 subunit that interacts with both SMC2 and SMC4 neck domains and clamps the DNA from above (Fig. 3B). The rest of the SMC coiled coil and the hinge domain remain flexible and are unresolved in the density map. The CAP-H2 N-terminal domain is bound to SMC2 neck domain further encircling the DNA molecule, in a conformation that resembles the “close gate” observed in other SMC complexes bound to DNA (Fig. S5H-J). Despite being present during sample preparation, no EM density was obtained for the CAP-G2 subunit. Accordingly, M18BP1^P^, that binds to condensin II through the CAP-G2 subunit, is also not visible in the final density map. This phenomenon is reminiscent of what is observed in structures of yeast condensin bound to the DNA (*28-30*), where Ycg1, the yeast CAP-G2 homologue, is also found only flexibly connected to the head module.

### Cryo-EM structure of condensin II in the autoinhibited state

While attempting to solve the cryo-EM structure of a condensin II-M18BP1^P^ bound to DNA in presence of ADP-Beryllium Fluoride (ADP-BeF_3_), most particles aggregated on the grid and were unusable for image processing (Fig. S6A). However, we identified a subpopulation of individual particles lacking both M18BP1^P^ and DNA. The resulting structure of condensin II bound to two ADP-BeF_3_ molecules was refined at an average resolution of 4.2 Å (Fig. 3C, Fig. S6A-G). The SMC2 and SMC4 ATPase heads are engaged and adopt a conformation competent for ATP hydrolysis with the two coiled coils of SMC2 and SMC4 that separate at the neck domains but reconnect after ca. 25 residues, forming a parallel coiled coil structure (Fig. 3D, Fig. S6F,H). The elbow and hinge domains are only visible at low contour level due to intrinsic flexibility (Fig. S6E). The CAP-G2 subunit hooks both SMC subunits by binding simultaneously to the SMC2 head and to the SMC4 coiled coil region. Additionally, the CAP-H2 N-terminal domain is bound between the CAP-G2 subunit and the SMC2 ATPase head and undergoes a substantial structural rearrangement when compared to the DNA-bound condensin II structure (Fig. 3D, Fig. S6I,J). The CAP-H2 C-terminal domain binds the SMC ATPase heads in a conformation reminiscent of other SMC structures (*28, 29, 31*), while its long unstructured domain interacts with both HEAT repeats (residues 148-202 bound to CAP-D3 and residues 316-367 bound to CAP-G2) (Fig. S7A). Finally, the CAP-D3 subunit is bound to CAP-G2 via its HEAT docker helices and remains on the edge of the complex without interacting with the SMC moieties (Fig. 3C,D). Additional CAP-D3 C-terminal residues (1454-1465 and 1469-1476), could be fitted and refined into the density map after modelling with AlphaFold3 (*32*) (Fig. S7B,C). Interestingly, this region of the CAP-D3 C-terminal tail has been shown to mediate an auto-inhibitory interaction with the CAP-G2 subunit (*20*). In agreement with this observation, the cryo-EM structure reveals that the CAP-D3 C-terminal tail anchors the N-terminal domain of CAP-H2 and the CAP-G2 subunit (around residues 386 – 418), stabilizing this tripartite interaction (Fig. S7B,C). Thus, the cryo-EM structure of condensin II in presence of ADP-BeF_3_ reveals a novel conformation characterized by an unusual arrangement of HEAT repeats-containing subunits that leads to an autoinhibited conformation of the condensin II complex, as highlighted by the superimposition with the structure of the DNA-bound condensin II–M18BP1^P^ complex (Fig. S7D).

### M18BP1^P^ relieves condensin II autoinhibition

To address how M18BP1^P^ activates condensin II, we obtained a novel cryo-EM structure of the condensin II-M18BP1^P^ complex at a higher resolution (3.8 Å local resolution) (Fig. 4A, Fig. S8). While the overall architecture closely resembles the previously published condensin II–M18BP1^P^ complex (*18*), the higher-resolution map reveals additional density for M18BP1^P^ on the surface of the CAP-G2 subunit (Fig. 4A,C). Comparison between the structure of condensin II in its autoinhibited conformation with the structure of the condensin II-M18BP1^P^ complex highlights the dramatic rearrangement of CAP-G2 and CAP-D3 subunits, while the overall conformation of the SMC2, SMC4 and CAP-H2 subunits remains largely unchanged. In the presence of M18BP1^P^, the two HEAT repeats-containing subunits act as a rigid body, undergoing a roto-translational movement spanning ca. 60 angstroms (Fig. 4B,C). This movement disrupts the contacts between CAP-G2 and both the SMC2 ATPase head and the SMC4 coiled coil. Importantly, the novel SMC2 head/CAP-G2 interface observed in the apo structure is in close proximity of the M18BP1^P^ binding site for CAP-G2 (Fig. 4C). An AlphaFold3 prediction validates the extended interaction interface between M18BP1^P^ and CAP-G2 observed in the cryo-EM map, including M18BP1 residues whose phosphorylation enhance M18BP1 binding to condensin II (Fig. S7E,F) (*18*). This suggests that M18BP1 binding to the CAP-G2 subunit could destabilize the CAP-G2/SMC2 interaction, promoting the structural rearrangement of the HEAT repeats-containing subunits. To test this hypothesis, we performed microscale thermophoresis (MST) experiments to directly assess the interaction between the SMC2/4 dimer and CAP-G2. Titration of SMC2/4 against a fixed concentration of fluorescently labelled CAP-G2 resulted in a measurable affinity (dissociation constant K_*D*_ equal to 1.3 +/-0.6 μM) (Fig. 4D). In contrast, no binding was detected in the presence of a 1-fold excess of M18BP1^P^, demonstrating that M18BP1^P^ binding prevents the interaction between the SMC2/4 dimer and CAP-G2 (Fig. 4D). This observation, together with the structural rearrangements observed in the condensin II - M18BP1^P^ complex in isolation and when bound to the DNA, suggests that M18BP1^P^ binding facilitates the detachment of CAP-G2 from its autoinhibitory position against the SMC2 ATPase head, thereby enabling the transition of condensin II into the DNA-bound clamped conformation (Fig. 4E).

**Fig. 4.**
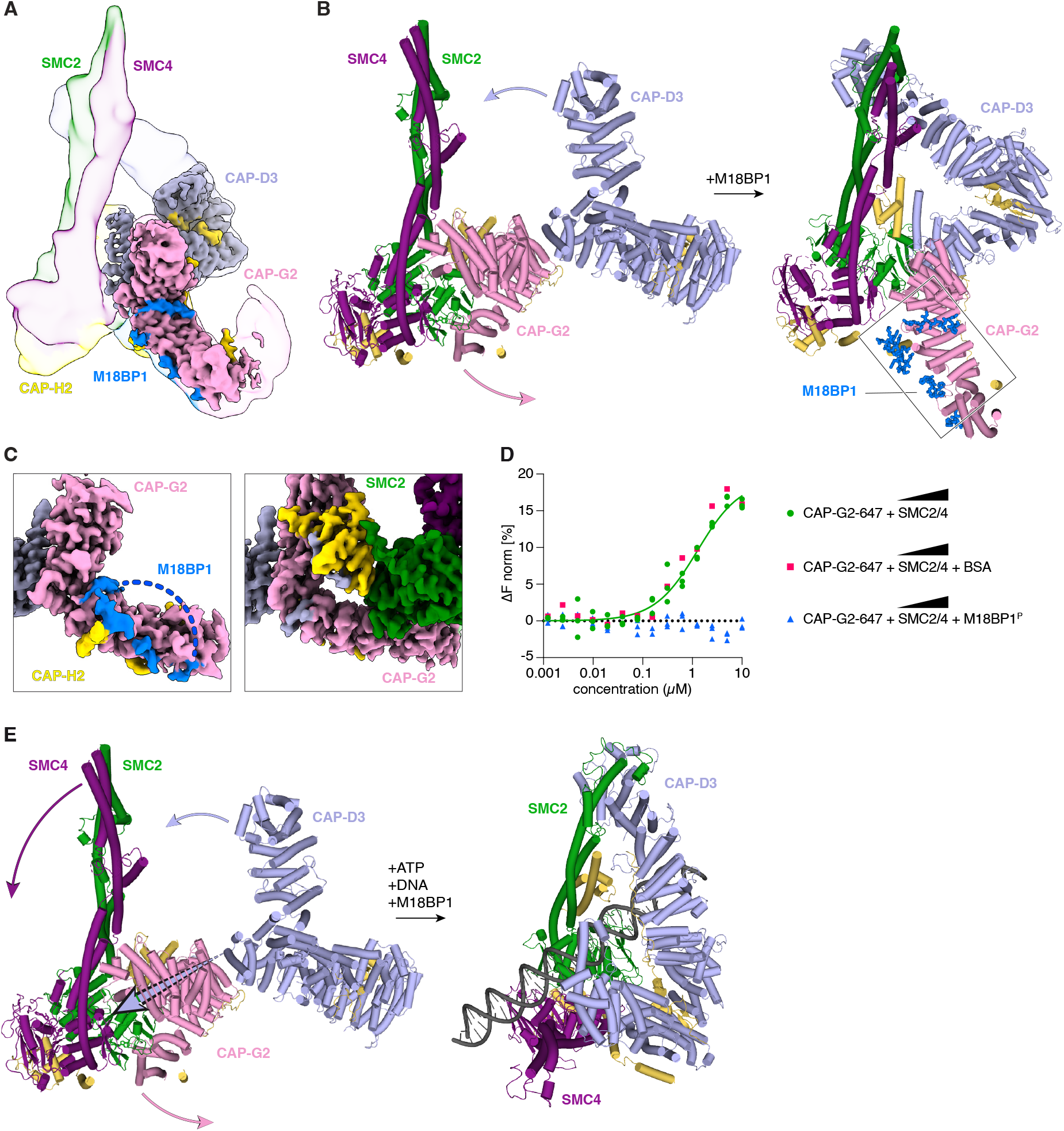
M18BP1^P^ binding relieves an autoinhibitory conformation of condensin II. (**A**) Cryo-EM map of condensin II bound to M18BP1^P^. (**B**) Model of condensin II bound to two ADP-BeF_3_ molecules with detail on the movement of the CAP-G2 and CAP-D3 subunits in the transition to the M18BP1^P^ bound condensin II on the right. Arrows indicate the roto-translational movement of the heat repeats. (**C**) Comparison between the CAP-G2 interaction surface in presence (left) and absence of M18BP1^P^ (right). (**D**) Microscale thermophoresis curves with titration of SMC2/4 from 10 µM to 240 pM over a fixed concentration (10 nM) of CAP-G2-647. Blue data points represent experiments performed in presence of 2 µM M18BP1^P^. Red data points represent experiment performed in presence of 2µM BSA. The curve fit was obtained by combining the data from three independent experiments. (**E**) Model of condensin II bound to two ADP-BeF_3_ molecules with arrows representing the movement of the SMC4, CAP-G2 and CAP-D3 subunits in the transition to the DNA bound condensin II on the right.

## Discussion

The activation of condensin II at mitotic onset is a multilayered process that involves specific phosphorylation of its subunits by mitotic kinases (*20, 33*) and the simultaneous binding of the activator M18BP1 protein, which outcompetes the inhibitory MCPH1 protein, a process also regulated by phosphorylation. Here, we provide detailed snapshots of the molecular mechanism defining the activation of condensin II by M18BP1^P^ and the resulting effects on DNA compaction activity. Optical tweezers combined to fluorescence microscopy experiments reveal that binding of condensin II to DNA in presence of ATP is increased in both frequency and duration when M18BP1^P^ is present. Moreover, the presence of M18BP1^P^ favors the transition from a diffusive condensin II to a non-diffusive, stably anchored complex on DNA. These results align and explain previous in vivo evidence showing that M18BP1^P^ is essential for condensin II localization on chromatin (*18, 19*).

The cryo-EM structural data of DNA-bound and apo M18BP1^P^-condensin II complexes, provide a potential molecular mechanism for the change in behavior of condensin II observed with the optical tweezers experiments (Fig. 5). In absence of both M18BP1 and DNA, condensin II adopts a conformation whereby the CAP-G2 subunit bridges the ATPase heads of the SMC subunits, forming a closed architecture that represents an autoinhibited state not competent for DNA entrapment. Binding of M18BP1^P^ promotes an extensive remodeling of the condensin II architecture, displacing CAP-G2 from its inhibitory position and priming the complex for DNA engagement. This structural transition is more evident in the clamped DNA-bound conformation, solved in the presence of both ADP-BeF_3_ and M18BP1^P^, where CAP-G2 disengages from the core of the complex, likely remaining associated through the interaction with the flexible CAP-H2 linker. Importantly, phosphorylation of the CAP-D3 C-terminal tail has additionally been characterized to be important for releasing CAP-G2 from its autoinhibited state (*20*). In agreement with this evidence, the cryo-EM structure of the condensin II complex in its autoinhibited conformation reveals the intimate connection between both the CAP-D3 heat docker domain and the CAP-D3 C-terminal tail with the CAP-G2 subunit (Fig. 3C,D and Fig. S7B,C). In summary, the two autoinhibitory mechanisms provide a redundant safe-belt mechanism to ensure a stringent control of condensin II activation. Together, the binding of M18BP1^P^ to CAP-G2 and the phosphorylation of the CAP-D3 tail trigger the transition of condensin II into an active complex capable of clamping the DNA, with the CAP-G2 subunit disengaging from the core of the complex. Similarly in yeast, the condensin I homologue (and CAP-G2 orthologue) Ycg1 disengages from the core SMC complex upon DNA binding, but this occurs independently of auxiliary factors, highlighting how higher eukaryotes have evolved additional regulatory layers to fine-tune condensin II activation. This likely holds true for human condensin I, which exhibits a higher degree of conservation with yeast condensin and, additionally, does not require additional factors to compact DNA in the optical tweezers experiments (Fig. S3B). It is interesting to note, that the CAP-D3 tail is not present in the map of the condensin II-M18BP1^P^ complex, suggesting that M18BP1 binding could favor the dissociation of the CAP-D3 tail from the CAP-G2 subunit, thus overriding the need for phosphorylation of the CAP-D3 tail to activate condensin II. This hypothesis is in line with the evidence that M18BP1 is required and sufficient to trigger pre-mature chromosome condensation in absence of MCPH1 and phosphorylation of the CAP-D3 tail (*18*). Thus, M18BP1^P^ binding likely represents the major activating mechanism of condensin II at mitotic onset, while CAP-D3 C-terminal tail phosphorylation might additionally contribute by disfavoring re-binding of the CAP-G2 subunit in its autoinhibitory conformation.

**Fig. 5.**
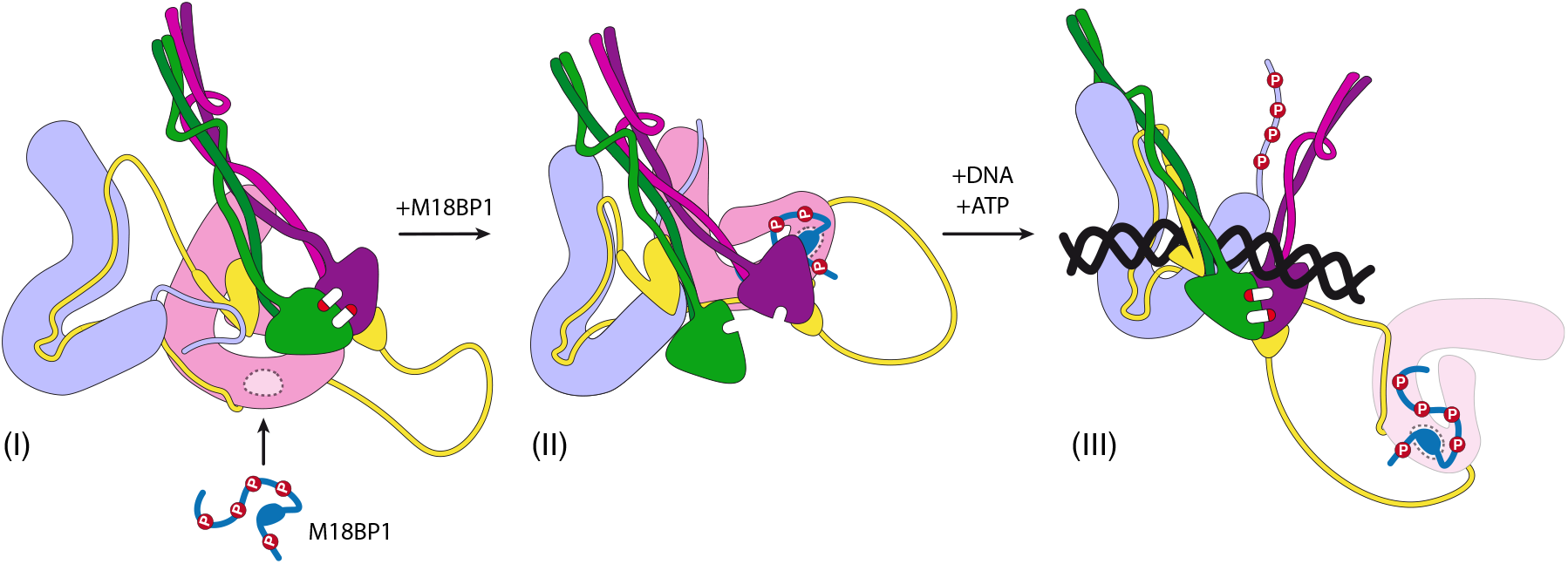
Model of condensin II activation. (I) Condensin II in the autoinhibited conformation, with CAP-G2 (pink) that bridges the ATPase heads. (II) The autoinhibited conformation of condensin II is released upon M18BP1^P^ binding. (III) Phosphorylation of the CAP-D3 tail, release of the CAP-G2 subunit, ATP mediated opening of the SMC2/4 coiled coil and DNA binding induce the formation of the DNA clamped conformation.

The activity of SMC complexes has been largely associated with the ability to form loops of DNA, as demonstrated by single molecule assays where individual complexes actively form loops of DNA in a process that is ATP dependent (*8, 12, 34-36*). The optical tweezers experiments shown here show how the formation of multi protein assemblies both in condensin I and condensin II play an important role in the compaction of the DNA molecule. This corroborates recent models that propose how loop extrusion by individual complexes is needed at the initial stages of mitotic chromosome folding, and is then replaced by another mechanism where neighboring non extruding condensins interact with each other to form functional assemblies (*9, 11, 15, 37*). Further work will be required to define the molecular determinants of the multiprotein assemblies observed here and to assess whether preventing their formation influences the architecture and stability of mitotic chromosomes.

## Acknowledgements

We thank the Cryo-Electron Microscopy and Biophysics Units of the National Facility for Structural Biology at Human Technopole, for technical support and assistance.

## Author contributions

Conceptualization: AB, AV

Methodology: AB, JA

Investigation: AB, SC, JA, VC

Visualization: AB

Funding acquisition: AV

Project administration: AB, AV

Supervision: AV

Writing – original draft: AB

Writing - revised draft; AB, AV

Writing – review & editing: AB, SC, JA, VC, AV

## Data, code, and materials availability

All data are available in the main text or the supplementary materials. The cryo-EM map and atomic coordinates of the condensin II complex in autoinhibited conformation have been deposited in the Electron Microscopy Data Bank (EMDB) and Protein Data Bank (PDB) with accession code EMD-56221 and 9TSZ. The cryo-EM map and atomic coordinates of the condensin II-DNA complex have been deposited in the Electron Microscopy Data Bank (EMDB) and Protein Data Bank (PDB) with accession code EMDB-56247 and 9TTP. The cryo-EM map and atomic coordinates of the condensin II-M18BP1 complex have been deposited in the Electron Microscopy Data Bank (EMDB) and Protein Data Bank (PDB) with accession code EMD-56273 and 9TUU.

## SUPPLEMENTARY Figure Legends

**Fig. S1. M18BP1 only associates with DNA in presence of condensin II**. (**A**,**B**) Schematic representation of the five-channel microfluidic device used to test protein binding to individual λ-DNA molecules. Gray arrow represents the different experimental phases. (**A**) M18BP1 labelled with AlexaFluor 488 (M18BP1-488) is introduced into the protein channel in the presence of ATP, without condensin II (CII). (**B**) M18PB1-488 is introduced together with condensin II labelled with LD655 (CII-655) in the presence of ATP. (**C**) Representative kymograph showing no detectable DNA binding of M18BP1-488 in the absence of condensin II. (**D**) Representative kymograph showing DNA binding of M18BP1-488 in the presence of condensin II -655 and ATP.

**Fig. S2. Quantification of fluorescence intensity traces from relax-stretch assay**. (**A**) Schematic representation of the five-channel microfluidic device used to test protein binding to individual λ-DNA molecules. Yellow arrow represents the different experimental phases. (**B**) Force-distance curve of experiment shown in Fig. 2C and in panel C. (**C**) Representative kymograph of relax–stretch assay performed with condensin II 2 nM and M18BP1^P^ 10 nM and ATP 3 mM. (**D**) Quantification of fluorescence intensity of traces from kymograph shown in panel A, reporting mean values and Gaussian fits. Blue histograms arise from traces before relaxation while green histograms arise from traces after relaxation and DNA compaction.

**Fig. S3. DNA compaction by condensin II is concentration dependent**. (**A**) In magenta, force– distance curves of λ-DNA molecule in presence of condensin II 1 nM, and ATP 3 mM. In Blue, force–distance curves of λ-DNA molecule after adding M18BP1P at 10 nM in the same channel of condensin II. (**B**) Force-distance curves of λ-DNA molecule in presence of condensin I 0.5 nM ATP 3 mM from five technical replicates. (**C**) Kymographs showing relax-stretch assay of single λ-DNA molecules in the presence of 488-labelled M18BP1^P^ at 20 nM and increasing concentrations of condensin II (CII): 2 nM, 1 nM, 0.5 nM, and 0.25 nM. White dotted lines outline beads position during the experiment. Composition of microfluidic protein channel is indicated schematically to the left of each kymograph. (**D**) Force-distance (FD) curves obtained with the relax-stretch assay in panel C. Curves represent DNA force-distance profiles following incubation with the respective condensin II concentrations. DNA compaction correlates with shortening of the DNA molecule, visible as a shift from the typical force-distance profile expected for a DNA molecule of given length (red line).

**Fig. S4. DNA compaction activity dependency on ATP hydrolysis and protein concentration**. (**A**,**B**,**C**) Force–distance curves of λ-DNA molecule in three different experimental conditions as described in figure panels: (**A**) condensin II^WT^ with M18BP1^P^ in presence of ATP. (**B**) condensin II^QL^ with M18BP1^P^ and ATP. (**C**) condensin II^WT^ with M18BP1^P^ and ADP-BeF_3_. High concentration (2 nM) and low concentration (0.25 nM) are shown in green and magenta, respectively. High concentration experiments were performed in triplicate for each condition, whereas low concentration experiments were repeated eight to thirteen times.

**Fig. S5. Cryo-EM data analysis of condensin II bound to DNA**. (**A**) Representative micrograph. (**B**) 2D class averages from full dataset. (**C**) Fourier Shell Correlation (FSC) curves for raw map obtained with RELION. (**D**) Model to map Fourier Shell Correlation (FSC) curves obtained with phenix. (**E**) Schematic representation of the cryo-EM data processing pipeline. (**F**) Model fit to map for the whole condensin II tetramer (SMC2 green, SMC4 purple, CAP-D3 lightblue and CAP-H2 yellow) bound to DNA (black). (**G**) Orientation distribution in final set of refined particles. (**H**) Model of the CAP-H2 N-terminal domain (yellow), in the conformation of human condensin II in the clamped DNA conformation. (**I**) Model of Rad21 N-terminal domain (yellow), in the conformation of human cohesin in the clamped DNA conformation (PDB: 6WG3). (**J**) Model of Brn1 N-terminal domain (yellow), in the conformation of Yeast condensin in the clamped DNA conformation (PDB: 7Q2X).

**Fig. S6. Cryo-EM data analysis of condensin II bound to two ADP-BeF**_**3**_ **molecules**. (**A**) Representative micrograph. (**B**) 2D class averages from full dataset. (**C**) Fourier Shell Correlation (FSC) curves for raw map obtained with RELION. (**D**) Model to map Fourier Shell Correlation (FSC) curves obtained with phenix. (**E**) Schematic representation of the cryo-EM data processing pipeline. (**F**) Model fit to map for the whole condensin II pentamer (SMC2 green, SMC4 purple, CAP-D3 lightblue, CAP-G2 pink, CAP-H2 yellow), with zoom in onto the regions with highest resolution. (**G**) Orientation distribution in final set of refined particles. (**H**) Detail of the ABC ATPase domain with the two ADP-BeF_3_ molecules represented as sticks. (**I**) Detail of the structure and position of the CAP-H2 N-terminal domain sandwiched between the SMC2 neck and the CAP-G2 subunit. (**J**) Upper panel, detail of the CAP-H2 N-terminal domain sandwiched between the SMC2 neck and CAP-G2 subunit in the structure of condensin bound to two ADP-BeF_3_ molecules. Lower panel, detail of CAP-H2 N-terminal domain rearranged in the condensin II DNA bound conformation.

**Fig. S7. Condensin II in the inhibited conformation**. (**A**) Detail of the CAP-H2 interaction sites with condensin II subunits. (**B**) Cryo-EM density map of condensin II bound to two ADP-BeF_3_ molecules zoomed in the region of interaction of the CAP-D3 C-terminal tail with CAP-G2 and the CAP-H2 N-terminal helices. (**C**) Model of condensin II bound to two ADP-BeF_3_ molecules, with highlighted residues of CAP-D3 tail that interact with CAP-G2 and the CAP-H2 N-terminal helices. (**D**) Superimposition of condensin II structure in the autoinhibited conformation with condensin II structure bound to DNA colored in grey. Dashed line represents potential clash between the SMC4 subunit of condensin II and the DNA molecule. (**E**) Alphafold3 prediction of the CAP-G2 - M18BP1 interaction, with relative PAE plot in panel (**F**).

**Fig. S8. Cryo-EM data analysis of condensin II bound to M18BP1. (A)** Representative micrograph. (**B**) 2D class averages from full dataset. (**C**) Schematic representation of the cryo-EM data processing pipeline. (**D**) Fourier Shell Correlation (FSC) curves for raw map obtained with RELION. (**E**) Model to map Fourier Shell Correlation (FSC) curves obtained with phenix. (**F**) Model fit to map for the whole condensin II - M18BP1 complex (SMC2 green, SMC4 purple, CAP-D3 lightblue, CAP-G2 pink, CAP-H2 yellow, M18BP1 blue), with zoom in onto the regions with highest resolution. (**G**) Orientation distribution in final set of refined particles.

**Fig. S9. Condensin II and M18BP1 labelling**. (**A**) Size exclusion profile (Superose 6 increase, 24 mL volume) of condensin II following labelling reaction with LD655 - CoA and SFP transferase. In grey and red respectively, absorbance at 280 nm and 650 nm. (**B**) SDS page gel of fractions from size exclusion in (A). Left panel, stained with Coomassie blue. Right panel, imaged by exciting at 650 nm. (**C**) Size exclusion profile (Superdex 200 increase, 2.4 mL volume) of M18BP1 following labelling reaction with AlexaFluor 488 - maleimide. In grey and blue respectively, absorbance at 280 nm and 495 nm. (**D)** SDS page gel of unlabelled (A) and labelled (B) M18BP1. Left panel, stained with Coomassie blue. Right panel, imaged by exciting AlexaFluor 488.

## Materials and Methods

### Protein expression

Human condensin II, condensin II sub-complexes, condensin II Q-loop mutant and condensin I were assembled into biGBac vectors as described previously (*12, 38, 39*). Recombinant bacmids were generated via Tn7 transposition in DH10EMBacY (Geneva Biotech) cells and transfected into Sf9 cells (Thermo Fisher Scientific, 11496015). Virus-containing supernatant was harvested after 3 days and further amplified in Sf9 cells for 3 additional days. For protein expression, amplified virus was used to infect High Five insect cells (Thermo Fisher Scientific, B85502). High Five insect cells were harvested after 3 days and centrifuged at 5,000 g for 30 minutes at 4°C and flash frozen in liquid nitrogen. MBP-M18BP1_873-1132_ and MBP-M18BP1_983-1045-7A_ were expressed in Rosetta (DE3) (MilliporeSigma (Novagen), 71401-4). Cells were grown in Terrific Broth media at 37°C. Upon reaching OD of about 4, cells were induced for 2 hrs at 30°C, then centrifuged at 5,000 g for 30 minutes at 4°C and flash frozen in liquid nitrogen.

### Protein purification

Cell pellets containing condensin II complexes were resuspended in 20 mM HEPES pH 8.0, 300 mM KCl, 5 mM MgCl_2_, 1 mM DTT, 10% glycerol, supplemented with 1 Pierce EDTA-free protease inhibitor tablet (Thermo Scientific) and 25 U/mL Benzonase (Sigma). Cells were lysed using a dounce homogenizer followed by sonication for 5 minutes (10 seconds on and 20 seconds off), and clarified by centrifugation for 40 minutes at 4°C using Beckman Coulter F20 rotor at 20k RPM. Clarified lysates were loaded onto a Strep-Tactin XT column (IBA Lifesciences), washed with lysis buffer, and eluted with the same buffer supplemented with 50 mM biotin (Sigma). Protein fractions were pooled and diluted 3-fold in Buffer A (20 mM HEPES pH 8.0, 5 mM MgCl_2_, 5% glycerol, 1 mM DTT). The diluted sample was loaded onto a HiTrap Heparin HP column (Cytiva), washed with Buffer A containing 250 mM NaCl, and eluted with Buffer A containing 500 mM NaCl. Finally, protein was further purified by size exclusion chromatography using Superose 6 Increase 16/600 (Cytiva) equilibrated in 20 mM HEPES pH 8, 300 mM KCl, 5 mM MgCl_2_, 1 mM DTT, 10% glycerol. Fractions containing purified complexes were pooled, concentrated and flash frozen in liquid nitrogen.

Bacterial pellets expressing MBP-M18BP1 constructs were resuspended in 30 mM HEPES pH 7.5, 200 mM NaCl, 10% glycerol, 2 mM DTT, and lysed by sonication on ice at 60% amplitude, 30 second on and 30 seconds off for a total of 5 minutes. After centrifugation for 30 minutes at 4°C using Beckman Coulter F20 rotor at 20k RPM the supernatant was injected on a HisTrap column and eluted with increasing concentration of imidazole. The HisTrap purification was followed by a MBP affinity purification and finally a size exclusion chromatography using Superdex 200 16/600 column (Cytiva). Phosphorylation of M18BP1 was obtained by treating the purified protein overnight at 4 °C with CDK1:Cyclin-B:CKS1 complexes in presence of 2 mM ATP and 10 mM MgCl_2_ Phosphorylation of M18BP1 was obtained by treating the purified protein overnight at 4 °Cas described in Huis in ‘t Veld P. J. *et al*. (*40*).

For human condensin I, pellets were resuspended in lysis buffer (25 mM HEPES, pH 8.0, 150 mM NaCl, 5 mM MgCl_2_, 1 mM DTT, and 10% glycerol), supplemented with a protease inhibitor cocktail (Calbiochem) and 25 U/mL Benzonase (Sigma). Cells were lysed using a Dounce homogenizer, followed by sonication for 5 minutes (15 s on/15 s off cycles), and the lysate was clarified by centrifugation for 60 minutes at 4 °C using a Beckman Coulter F20 rotor at 20,000 rpm. The clarified lysate was loaded onto a StrepTrap™ HP 5 ml column(Cytiva) equilibrated with lysis buffer, and proteins were eluted with the same buffer supplemented with 50 mM biotin (Sigma). Eluted protein fractions were pooled and applied to a HiTrap Heparin HP column (Cytiva) equilibrated with heparin buffer (25 mM HEPES, pH 8.0, 150 mM NaCl, 5% glycerol, and 1 mM DTT). Proteins were eluted using a linear NaCl gradient from 150 to 1000 mM. Fractions of the highest purity were pooled and loaded onto a Superose 6 Increase 10/300 column (Cytiva) equilibrated with gel filtration buffer (25 mM HEPES, pH 8.0, 150 mM NaCl, and 1 mM DTT). Selected fractions were concentrated, flash-frozen in liquid nitrogen, and stored at −80 °C.

### Protein labelling

Labelling of condensin II complexes was achieved by covalently attaching fluorescent dyes conjugated to Coenzyme-A (CoA) to the YBBR tag of SMC2 subunit. Briefly, YBBR tagged condensin II at 1 µM was incubated with Lumidyne 655 fluorophore conjugated to CoA (LD655-CoA, Lumydine Technologies) at 5 µM, in presence of 0.5 µM SFP (4’-phosphopantetheinyl transferase) for 1 hr at 4 °C in the dark. Subsequently, the reaction mixture was purified over a Superose 6 Increase 10/300 GL column equilibrated in 30 mM HEPES pH 7.5, 150 mM NaCl, 2 mM DTT, 0.05% (v/v) Tween20 and 5% glycerol. (Fig. S9A). 50 μL fractions were collected and analysed by SDS–PAGE using 4–12% NuPAGE Bis-Tris gels (Invitrogen) running in MOPS buffer at 200 V for 45 min. Imaging of the gel was performed on Chemidoc with optimal settings for imaging at 655 nm, after which the gel was stained with Instant-Blue Coomassie protein stain (Abcam) (Fig. S9B). Labelled condensin II fractions were pooled, aliquoted and flash frozen in liquid nitrogen at a concentration of 200 nM. Degree of labelling (DOL) was calculated with the following formula, yielding a labelling efficiency of 64%.

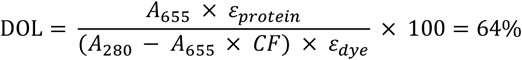

*A*_655_ = 0.131 nanodrop reading at 650 for LD655

*A*_280_ = 0.38 nanodrop reading at 280

*ε*_*protein*_ = 445210 *M*^−1^*cm*^−1^

*ε*_*dye*_ = 239000 *M*^−1^*cm*^−1^

*CF* = 0.03 *correction factor for Alexa Fluor* 655 *at* 280 *nm*

MBP-M18BP1_873-1132_ was labelled by covalently attaching AlexaFluor dyes to cysteine residues present on the protein. In order to remove DTT, MBP-M18BP1_873-1132_ at 12 µM was buffer-exchanged using ZebaSpin columns (0.5 mL volume) into 30 mM HEPES (pH 7.5), 150 mM NaCl, and 5% glycerol. Labelling of MBP-M18BP1_873-1132_ was achieved by covalently attaching AlexaFluor dyes to cysteine residues present in M18BP1. Following buffer exchange, the protein was incubated with AlexaFluor 488 C5 Maleimide at **xx** µM for 1 hr at 4 °C in the dark. The reaction was then quenched by addition of Tris buffer (pH 7.5) to a final concentration of 50 mM and subsequently loaded onto a Superdex 200 Increase 10/300 GL column equilibrated in 30 mM HEPES (pH 7.5), 150 mM NaCl, 2 mM DTT, 0.05% (v/v) Tween-20, and 5% glycerol. (Fig. S9C).50 μL fractions were collected and analyzed by SDS–PAGE using 4–12% NuPAGE Bis-Tris gels (Invitrogen) running in MES buffer at 200 V for 30 min. Imaging of the gel was performed on Chemidoc with optimal settings for imaging AlexaFluor 488, after which the gel was stained with Instant-Blue Coomassie protein stain (Abcam) (Fig. S9D)). Labelled MBP-M18BP1_873-1132_ fractions were pooled, aliquoted and flash frozen in liquid nitrogen at a concentration of 1 µM. Degree of labelling (DOL) calculated with previous formula using a correction factor of 0.11 (for AlexaFluor 488 at 280 nm), was 89%.

### Single molecule DNA binding experiments

Single-molecule DNA binding experiments were performed on a C-trap (LUMICKS) integrating optical tweezers, confocal fluorescence microscopy and microfluidics controlled by BlueLake software (LUMICKS). Every channel of the microfluidic device was first cleaned with 2% Helmanex, MQ water and then equilibrated in 20 mM HEPES (pH 7.5), 25 mM KCl, 5 mM MgCl_2_, 1 mM TCEP and 0.04 % (v/v) Tween20. Streptavidin-coated polystyrene beads, 0.005% w/v (4.35 μm, Spherotech), were injected into channel 1. Biotin-labelled λ-DNA molecules (about 2 pM) were flowed into channel 2. Channel 3 and 4 contained experimental buffer supplemented with 2 mM ATP. Channel 5 contained labelled protein diluted in experimental buffer. The optical trap was calibrated to achieve a trap stiffness of 0.2– 0.3 pN nm^−1^. After trapping two beads in channel 1, the DNA was captured in channel 2. Successful tether formation was verified in channel 3 by acquiring force-distance profiles, by extending the distance between the beads to 16.5 µm. The DNA was then transferred in the protein channel (channel 5) and the proteins were allowed to incubate for 1 minute at tension below 1pN. The incubation step was carried out in static conditions with all channel valves closed. During the incubation step, confocal fluorescence imaging was performed with laser excitation at 488, or 638 nm for AlexaFluor488 or LD655, respectively. Emission was collected through band-pass filters at 512/25 nm (blue) and a long-pass filter at 640 nm (red). Kymographs were generated by line-scanning along the DNA axis with a dwell time of 0.2 ms per pixel. All measurements were conducted at room temperature (21 °C). Five to six technical replicates were acquired per experimental condition.

For condensin I, experiments were carried out in the same manner, with slight changes in buffer composition that were previously optimized for condensin I stability. In this case, the microfluidic device was equilibrated with buffer containing 50 mM HEPES (pH 7.0), 50 mM NaCl, 1 mM MgCl_2_, 1 mM DTT, and 0.02% (v/v) Tween-20.

### Kymograph analysis and extraction of single-molecule trajectories

Kymographs were analyzed in Python using custom Jupyter Notebooks. Kymograph files were imported via the *lumicks*.*pylake* library and processed with the `KymoWidgetGreedy` tool to identify individual fluorescent trajectories. Traces were detected automatically using the greedy line-tracking algorithm (`KymoWidgetGreedy`), with the following parameters: pixel intensity threshold = 0.5, Gaussian smoothing sigma = 0.4, window size = 6 pixels, minimum track length = 8 pixels, maximum display intensity = 8, and origin correction enabled (`correct_origin=True`). Manual correction of the identified traces was performed to remove false positive and to connect traces when needed.

### Mean squared displacement (MSD) analysis

For each detected trace, the position along the kymograph (in µm) was extracted as a function of frame index and converted to a time point using the known temporal calibration of the acquisition (0.3 s per pixel). Traces shorter than three points or with insufficient temporal coverage were discarded. The mean squared displacement (MSD) was calculated in one dimension for each trajectory using a standard time-averaging approach. Briefly, for a given trace with positions *x*(*t*), the MSD at a lag time *τ*was computed as

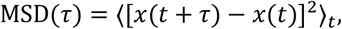

where the average is taken over all valid time points *t* within the trace. Lag times were sampled from 1 frame up to half the trajectory length to ensure sufficient statistics at longer lags. For each trace, the MSD as a function of lag time was saved to comma-separated values (CSV) files (^*^_msd.csv) together with the corresponding trace index and lag times.

To compare multiple kymographs and biological repeats, MSD curves from all traces and experiments belonging to the same condition were pooled. This was done by reading all ^*^_msd.csv files in condition-specific folders (e.g. for different nucleotide or protein conditions) and plotting the MSD versus lag time for each individual trace. This provided a visual assessment of the spread and shape of MSD curves across all trajectories and conditions.

### Estimation of diffusion coefficients

Diffusion coefficients were extracted from the initial, approximately linear regime of the MSD– lag time curves, assuming one-dimensional Brownian diffusion. For each trajectory, the MSD versus lag time data were first filtered to remove non-positive lag times and NaN values and sorted by increasing lag time. Only traces with at least three lag-time points and a total duration of at least 1 s (maximum lag time ≥ 1 s) were retained for fitting. The first *k* points (with *k* between 4 and 6, depending on trace length) were then used to perform an ordinary least-squares linear regression with an intercept,

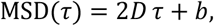

yielding a slope and intercept for each trace. The one-dimensional diffusion coefficient *D* was obtained as half of the fitted slope (*D* = slope/2). For each experimental condition, the per-trace diffusion coefficients were compiled into a single table and exported as a CSV file. The number of traces, mean diffusion coefficient, standard deviation, and standard error of the mean were computed per condition. To visualize the distribution of diffusion coefficients across conditions, violin plots were generated using Prism. These plots allowed direct comparison of the distribution and central tendency of diffusion coefficients obtained from multiple kymographs and independent experiments under the different biochemical conditions.

### Single molecule relax-stretch experiments

Single molecule relax-stretch experiments were performed under the same conditions as the DNA binding assays, except for the protein binding and relax-stretch procedure. A custom script was used to automate the experimental sequence. At the start of each measurement, all channel valves were closed, and the presence of a single DNA tether was verified in channel 3 by recording a force-distance profile. The tether was then transferred into channel 5 for protein binding, where it was held at a bead-to-bead distance of 14 µm under a force of less than 1 pN and incubated for 3 to 5 s. The tether was subsequently moved to channel 4 and an additional 5 s waiting period was included in this channel to confirm the presence of bound proteins. Protein activity was then allowed by decreasing the bead-to-bead distance to 5 µm and incubating for 1 min, after which the distance was increased to 17 µm. A speed of 1 µms^-1^ and 0.5 µms^-1^ were used to respectively decrease and increase the distance between the beads during the stretch-relax assay. Throughout the entire procedure, the applied force was continuously monitored, and confocal fluorescence imaging was conducted under the same conditions as described for the single molecule DNA binding experiments.

### Analysis of Force - Time and Force - Distance data

Force data were extracted directly from C-Trap .*h5* files using the *lumicks*.*pylake* Python library. For each dataset, the force channel (`Force LF / Force 2x`) was used to export the force values from the time window of the stretch-relax assay. Time (seconds) and force values (pN) were retrieved from the . `seconds` and . `data` attributes, respectively. The force–time trace was exported and plotted using GraphPad Prism, with fixed axis limits and a 0-pN reference line for consistent visualization.

Force–distance curves were obtained by reading the corresponding force (`Force LF / Force 2x`) and distance (`Distance / Distance 1`) channels from each .*h5* dataset over a defined time window. These values were exported to a CSV file and FD curves were plotted using GraphPad Prism, with force on the y-axis and distance on the x-axis.

### Cryo-EM sample preparation

Sample for the structure of condensin II bound to the DNA was obtained by mixing purified condensin II at 0.5 µM and M18BP1_873-1132_ (previously treated with CDK1:Cyclin-B:CKS1) at 1 µM in presence of 0.5 mM ADP, 0.5 mM BeSO_4_ and 5 mM NaF and 235 bp DNA molecule (Table S2) at 2 µM in 30 mM HEPES pH 7.5, 25 mM NaCl, 5mM MgCl_2_, 2mM DTT, 0.03% beta-glucopyranoside and 0.05% Tween20. After 10 minutes incubation, sulfo-NHS-diazirine (sulfo-SDA, Thermo Scientific) at 1 mM was added to the mixture and incubated for 30 minutes at 25°C to allow the NHS moiety to react with lysine residues. The crosslinked protein material was then treated with UV light (waveleght 330-370 nm) on ice for 10 minutes. The crosslinking reaction was quenched with 30 mM TRIS pH 7.5, and the freshly prepared sample was applied to Quantifoil R1.2/1.3 300-mesh copper holey carbon grids, previously glow-discharged using a GloQube (Quorum) set at 30 mA for 60 seconds. Grids were blotted for 3 s under 100% humidity at 4°C before being plunged into liquid ethane using a Vitrobot Mark IV (Thermo Scientific).

Identical procedure was followed to prepare the sample for the structure of condensin II in the autoinhibited conformation, with the only difference that condensin II was added at 1 µM instead of 0.5 µM before crosslinking.

Sample for the structure of condensin II – M18BP1^P^ complex, was obtained by mixing purified condensin II at 1 µM and M18BP1_873-1132_ (previously treated with CDK1:Cyclin-B:CKS1) at 3 µM in presence of 0.5 mM ADP, 0.5 mM BeSO_4_ and 5 mM NaF for 30 min in buffer containing 30 mM HEPES pH 7.5, 200 mM NaCl, 2mM DTT and 0.04% Tween20. The sample was then run on a glycerol (10-25%) and glutaraldehyde (0-0.1%) gradient. Ultracentrifugation with a Sw60 rotor set on a G-force of 29k was performed for 16hrs at 4°C. After the GraFix fixation, the complex was run on a Superose 6 Increase 10/300 GL column equilibrated in 30mM HEPES pH 7.5, 200 mM NaCl, 2mM DTT and 0.04% Tween20. The peak fractions were concentrated and applied to Quantifoil R1.2/1.3 300-mesh copper holey carbon grids, previously glow-discharged using a GloQube (Quorum) set at 30 mA for 60 seconds. Grids were blotted for 3.5 s under 100% humidity at 4°C before being plunged into liquid ethane using a Vitrobot Mark IV (Thermo Scientific).

### Cryo-EM data acquisition

Micrographs for condensin II - DNA structure were acquired as EER movies on a Glacios (Thermo Scientific) transmission electron microscope operated at 200 kV with a Falcon 4i (Thermo Scientific) direct electron detector and Selectris X (Thermo Scientific) energy filter set with slit width of 10 eV. EPU (Thermo Scientific) software was used for automated data collection following standard procedures. A magnification of 79,000x was used for imaging, yielding a pixel size of 1.598 Å on images. The defocus range was set from -0.5 μm to -2 μm. Each micrograph was dose-fractionated to 60 frames, with a total dose of about 60 e-/Å^2^.

Micrographs for condensin II in autoinhibited conformation were acquired as EER movies on a Titan Krios G4 (Thermo Scientific) transmission electron microscope operated at 300 kV with a Falcon 4i (Thermo Scientific) direct electron detector and Selectris X (Thermo Scientific) energy filter set with slit width of 10 eV. EPU (Thermo Scientific) software was used for automated data collection following standard procedures. A magnification of 130,000x was used for imaging, yielding a pixel size of 0.7 Å on images. The defocus range was set from -0.5 μm to -2 μm. Each micrograph was dose-fractionated to 60 frames, with a total dose of about 60 e-/Å^2^.

Micrographs for condensin II – M18BP1 structure were acquired as EER movies on a Titan Krios G4 (Thermo Scientific) transmission electron microscope operated at 300 kV with a Falcon 4i (Thermo Scientific) direct electron detector and Selectris X (Thermo Scientific) energy filter set with slit width of 10 eV. EPU (Thermo Scientific) software was used for automated data collection following standard procedures. A magnification of 165,000x was used for imaging, yielding a pixel size of 0.748 Å on images. The defocus range was set from -0.5 μm to -2 μm. Each micrograph was dose-fractionated to 60 frames, with a total dose of about 60 e-/Å^2^.

### Cryo-EM image processing

Details of processing pipelines and refinement statistics are shown in Fig. S5, S6, S8 and Table S1. All steps of image processing were performed using RELION-4 (*41, 42*). Motion correction was performed using the Relion’s own implementation of the MotionCorr2 program (*43*), and the CTF parameters of the micrographs were estimated using the CTFFIND program (*44*).. Initially, particle picking was performed by using the Laplacian-of-Gaussian blob detection method in RELION-4. Class averages representing projections in different orientations were selected from the initial reference-free 2D classification and further used for 3D initial model and 3D classification. Particles aligning in the best 3D classes were used for Topaz (*45*) training and autopicking. Extracted particles were binned 3 times and subjected to 2D classification and 3D classification. The selected classes from 3D classification were subjected to 3D auto refinement followed by Bayesian polishing and CTF refinement. When needed, particle subtraction followed by 3D classification and refinement were performed to increase the local resolution of the density maps.

### Microscale thermophoresis

Microscale thermophoresis (MST) experiments were performed using fluorescently labelled CAP-G2 (10 nM) as the ligand, and the SMC2/4 dimer as the titrant. CAP-G2 was labelled through the strep tag with a AlexaFluor647 fluorescently labelled streptavidin. A 16-point 1:2 serial dilution of SMC2/4 was prepared starting from 20 µM in buffer containing 30 mM HEPES pH 7.5, 150 mM KCl, 0.5 mM TCEP, and 0.05% (v/v) Tween-20. Samples were mixed 1:1 (v/v) with labelled CAP-G2, incubated for 10–15 min at room temperature, and loaded into premium-coated capillaries. Measurements were performed using high MST power on a Monolith instrument (NanoTemper Technologies). To assess the effect of M18BP1, parallel experiments were carried out in the presence of 2 µM M18BP1 or 2 µM BSA as a control. Data were analyzed using GraphPad Prism, and binding curves were fitted with a one-site specific binding model to determine dissociation constants (K_*D*_).

**Table S1.** Cryo-EM data collection, refinement, and validation statistics.

| **Data collection and processing** | Condensin II + ADP-BeF<sub>3</sub> | Condensin II + ADP-BeF<sub>3</sub> + DNA | Condensin II + M18BP1<sup>P</sup> |
| --- | --- | --- | --- |
| Magnification | 105,000 | 79,000 | 165,000 |
| Voltage (kV) | 300 | 200 | 300 |
| Electron exposure e-/Å<sup>2</sup> | 50 | 60 | 60 |
| Defocus range (μm) | 0.8 to 2.0 | 0.8 to 2.0 | 0.5 to 1.5 |
| Pixel size (Å) | 1.2 | 1.589 | 0.748 |
| Symmetry imposed | C1 | C1 | C1 |
| Initial particle images (no) | 3.7 M | 1.2 M | 388 k |
| Final particle images (no) | 49k | 70k | 52 k |
| Map resolution (Å) | 4.3 | 6.7 | 4.1 |
| FSC threshold | 1.43 | 1.43 | 1.43 |
| Map resolution range (Å) | 4 to ~ 7 | 6 to ~ 10 | 3.8 to ~ 6.5 |
| **Refinement** |  |  |  |
| Initial model used | 9F5W | AlphaFold monomer predictions | 9F5W |
| Model resolution (Å) | 3.9 | 6.2 | 4.1 |
| FSC threshold | 0.143 | 0.143 | 0.143 |
| Map sharpening B factor (Å<sup>2</sup>) |  |  |  |
| **Model comparison** |  |  |  |
| Nonhydrogen atoms | 23364 | 21245 | 27267 |
| Protein residues | 2891 | 2421 | 3384 |
| **r.m.s deviations** |  |  |  |
| Bond lengths (Å<sup>2</sup>) | 0.005 | 0.005 | 0.004 |
| Bond angles (°) | 1.043 | 1.050 | 0.957 |
| **Validation** |  |  |  |
| MolProbity score | 1.64 | 1.58 | 1.42 |
| Clashscore | 8.27 | 6.58 | 5.66 |
| Poor rotamers (%) | 1.77 | 1.48 | 1.34 |
| **Ramachandran plot** |  |  |  |
| Favored (%) | 96.85 | 96.58 | 97.42 |
| Allowed (%) | 3.12 | 3.3 | 2.55 |
| Outliers (%) | 0.04 | 0.13 | 0.03 |

**Table S2.**
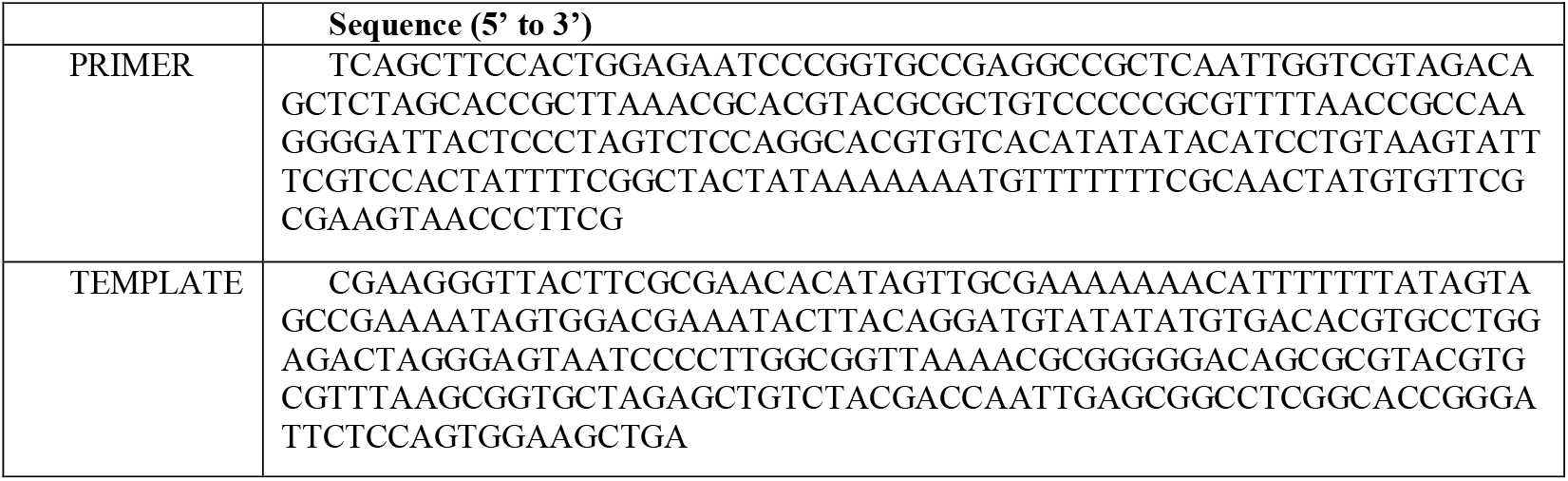
DNA sequence used for cryo-EM sample preparation.

