## Supplementary Figures for "Cryo-EM and single molecule visualization unravel the role of human condensin II activation by M18BP1 in driving DNA compaction"

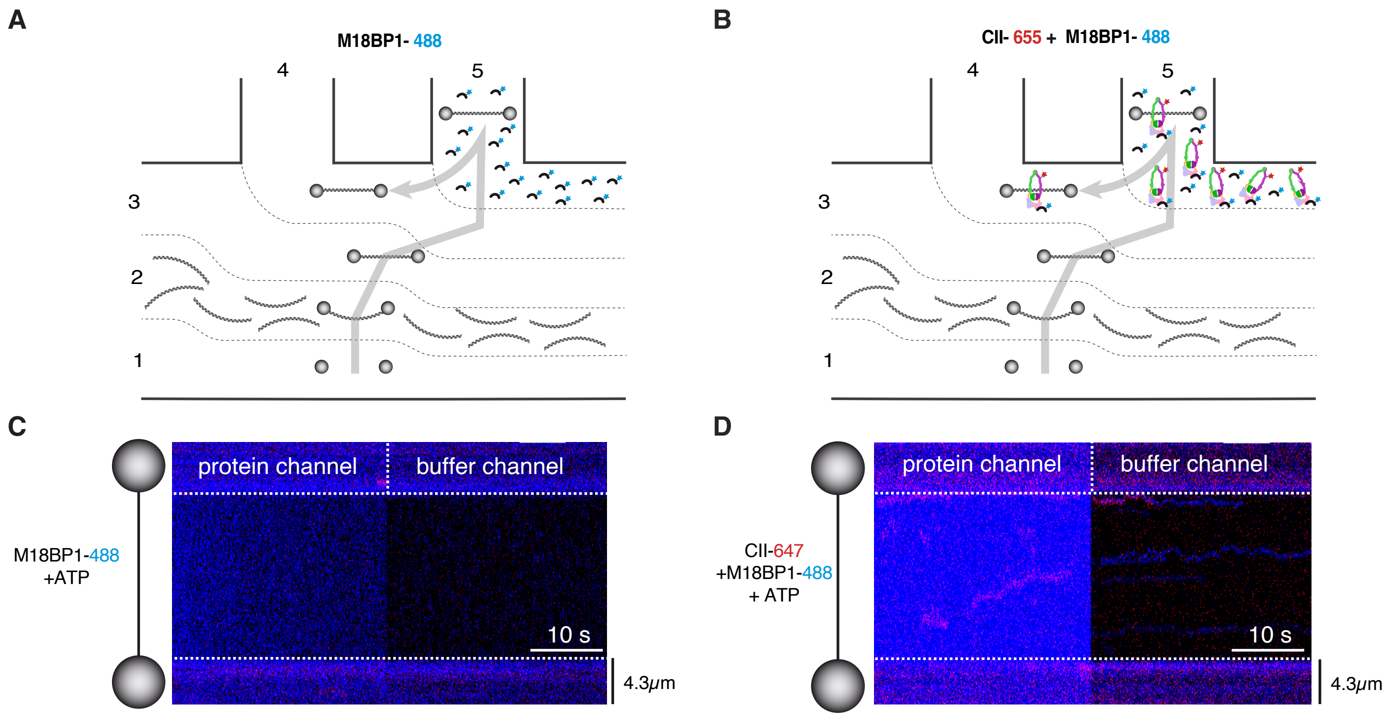


**Fig. S1. M18BP1 only associates with DNA in presence of condensin II.** (**A,B**) Schematic representation of the five-channel microfluidic device used to test protein binding to individual λ-DNA molecules. Gray arrow represents the different experimental phases. (**A**) M18BP1 labelled with Alexa Fluor 488 (M18BP1-488) is introduced into the protein channel in the presence of ATP, without condensin II. (**B**) M18BP1-488 is introduced together with condensin II labelled with LD655 (CII-655) in the presence of ATP. (**C**) Representative kymograph showing no detectable DNA binding of M18BP1-488 in the absence of condensin II. (**D**) Representative kymograph showing DNA binding of M18BP1-488 in the presence of condensin II-655 and ATP.

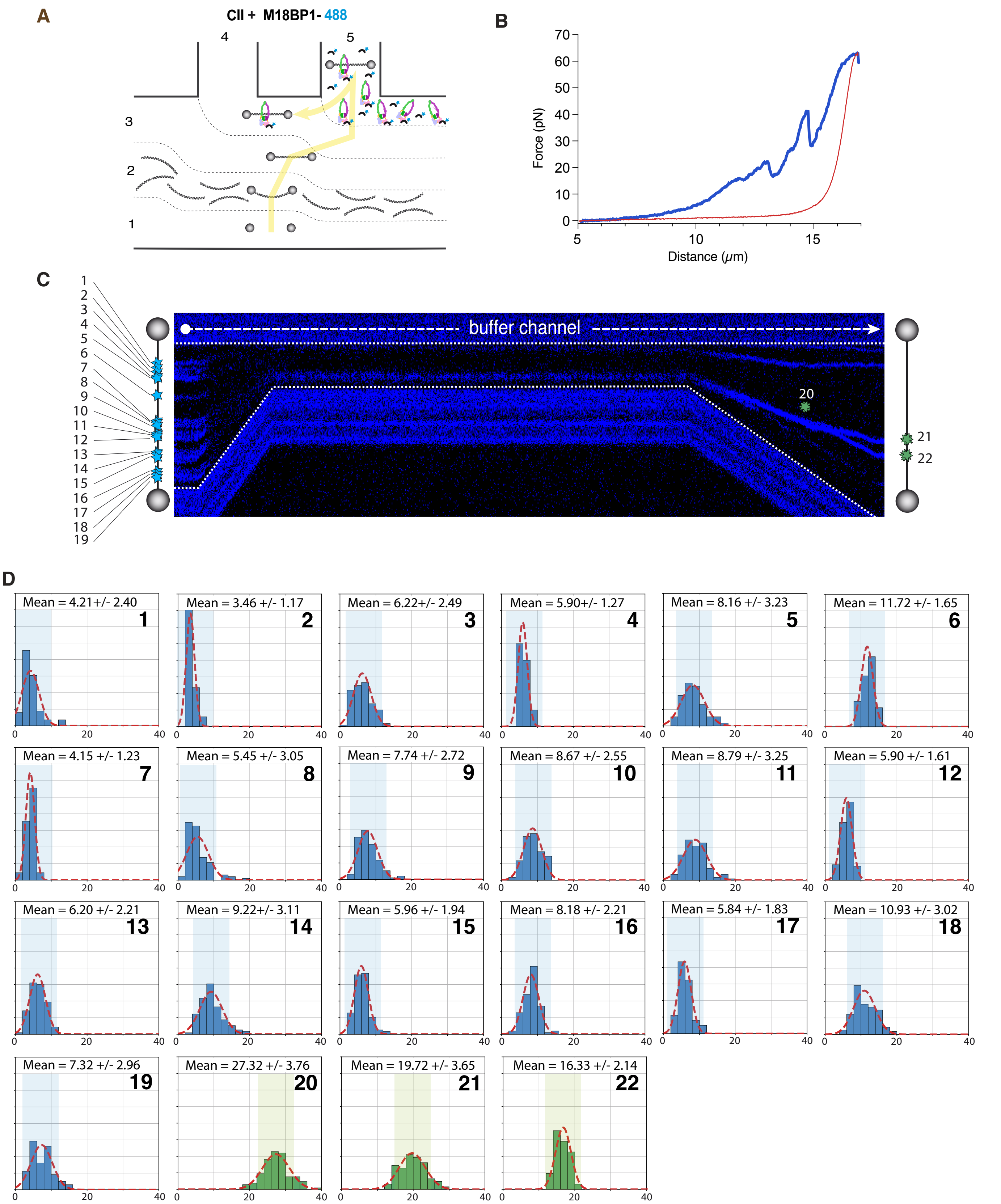


**Fig. S2. Quantification of flurescence intensity traces from relax-stretch assay.** (**A**) Schematic representation of the five-channel microfluidic device used to test protein binding to individual λ-DNA molecules. Yellow arrow represents the different experimental phases. (**B**) Force-distance curve of experiment shown in Fig. 2C and in panel C. (**C**) Representative kymograph of relax–stretch assay performed with condensin II 2 nM and M18BP1$^{P}$ 10 nM and ATP 3 mM. (**D**) Quantification of fluorescence intensity of traces from kymograph shown in panel A, reporting mean values and Gaussian fits. Blue histograms arise from traces before relaxation while green histograms arise from traces after relaxation and DNA compaction.

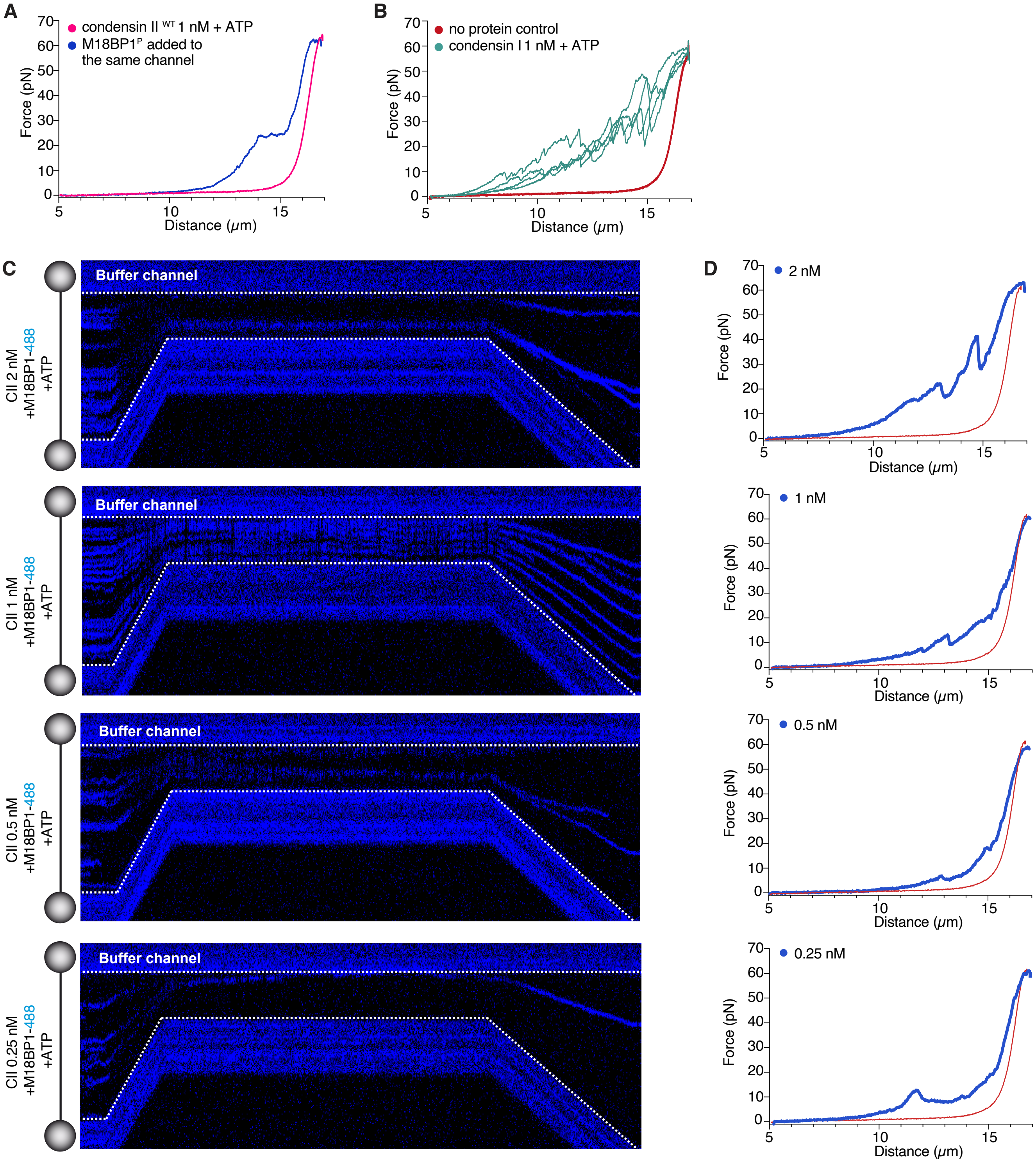


**Fig. S3. DNA compaction by condensin II is concentration dependent.** (**A**) In magenta, force–distance curves of λ-DNA molecule in presence of condensin II 1 nM, and ATP 3 mM. In Blue, force-distance curves of λ-DNA molecule after adding M18BP1$^{P}$ at 10 nM in the same channel of condensin II. (**B**) Force–distance curves of λ-DNA molecule in presence of condensin I 0.5 nM and ATP 3mM from 5 technical replicates. (**C**) Kymographs showing relax-stretch assay of single λ-DNA molecules in the presence of 488-labelled M18BP1$^{P}$ at 20 nM and increasing concentrations of condensin II (CII): 2 nM, 1 nM, 0.5 nM, and 0.25 nM. White dotted lines outline beads position during the experiment. Composition of microfluidic protein channel is indicated schematically to the left of each kymograph. (**D**) Force-distance (FD) curves obtained with the relax-stretch assay in panel C. Curves represent DNA force-distance profiles following incubation with the respective condensin II concentrations. DNA compaction correlates with shortening of the DNA molecule, visible as a shift from the typical force-distance profile expected for a DNA molecule of given length (red line).

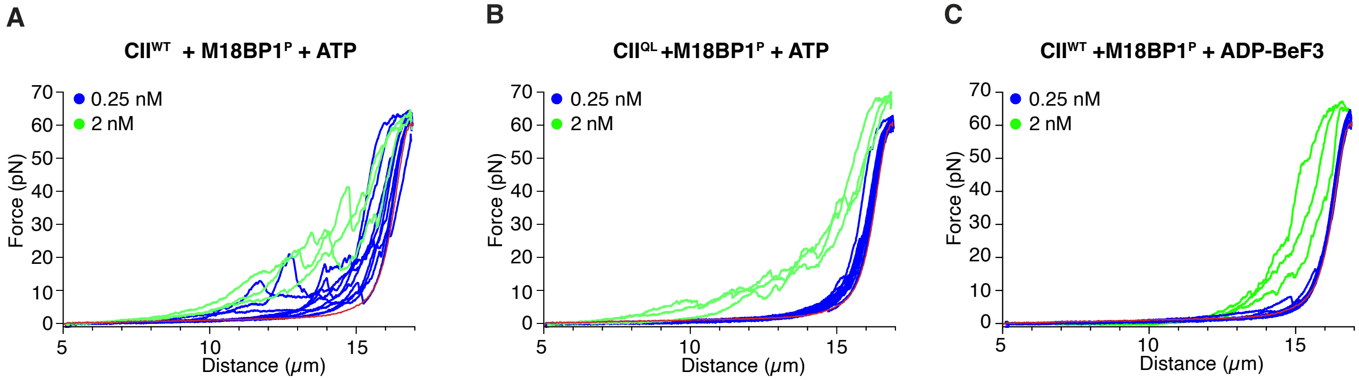


**Fig. S4. DNA compaction activity dependency on ATP hydrolysis and protein concentration.** (**A**,**B**,**C**) Force–distance curves of λ-DNA molecule in three different experimental conditions as described in figure panels: (**A**) condensin II$^{WT}$ with M18BP1$^{P}$ in presence of ATP. (**B**) condensin II$^{QL}$ with M18BP1$^{P}$ and ATP. (**C**) condensin II$^{WT}$ with M18BP1$^{P}$ and ADP-BeF$_3$. High concentration (2 nM) and low concentration (0.25 nM) are shown in green and blue, respectively. High concentration experiments were performed in triplicate for each condition, whereas low concentration experiments were repeated eight to thirteen times.

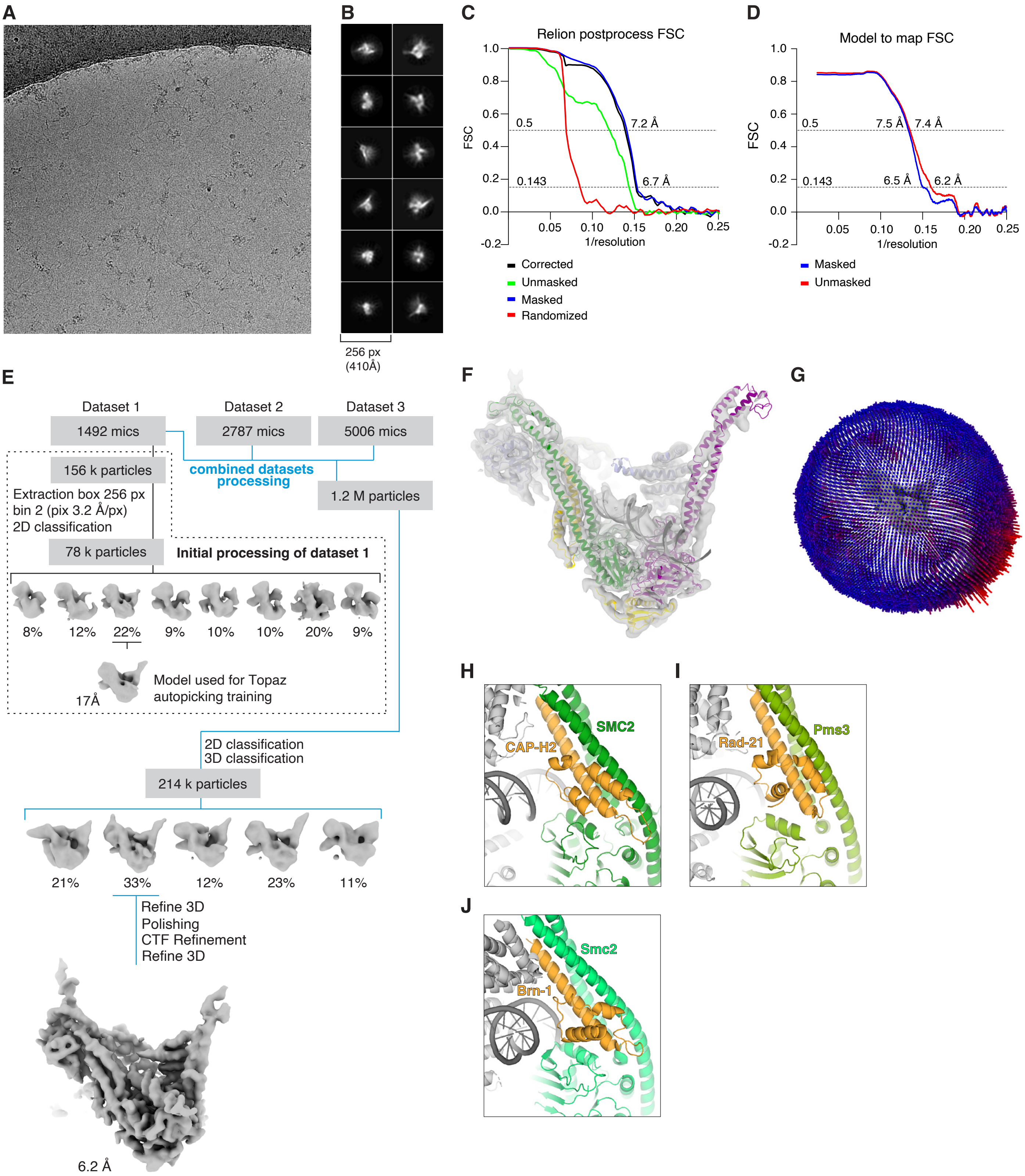


**Fig. S5. Cryo-EM data analysis of condensin II bound to DNA.** (**A**) Representative micrograph. (**B**) 2D class averages from full dataset. (**C**) Fourier Shell Correlation (FSC) curves for raw map obtained with RELION. (**D**) Model to map Fourier Shell Correlation (FSC) curves obtained with phenix. (**E**) Schematic representation of the cryo-EM data processing pipeline. (**F**) Model fit to map for the whole condensin II tetramer (SMC2 green, SMC4 purple, CAP-D3 lightblue and CAP-H2 yellow) bound to DNA (black). (**G**) Orientation distribution in final set of refined particles. (**H**) Model of the CAP-H2 N-terminal domain (yellow), in the conformation of human condensin II in the clamped DNA conformation. (**I**) Model of Rad21 N-terminal domain (yellow), in the conformation of human cohesin in the clamped DNA conformation (PDB: 6WG3). (**J**) Model of Brn1 N-terminal domain (yellow), in the conformation of Yeast condensin in the clamped DNA conformation (PDB: 7Q2X).

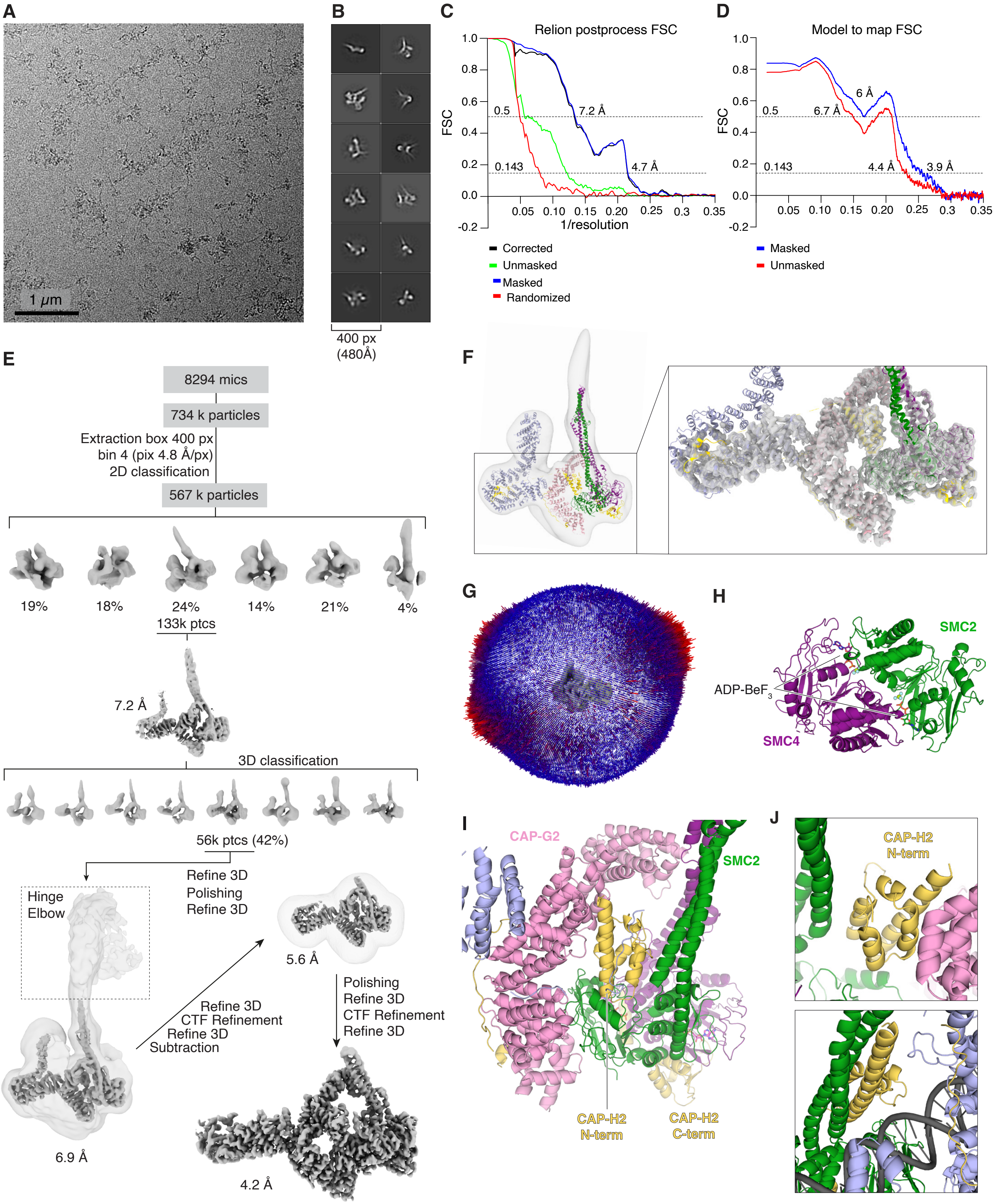


**Fig. S6. CryoEM data analysis of condensin II bound to two ADP-BeF$_3$ molecules.** (**A**) Representative micrograph. (**B**) 2D class averages from full dataset. (**C**) Fourier Shell Correlation (FSC) curves for raw map obtained with RELION. (**D**) Model to map Fourier Shell Correlation (FSC) curves obtained with phenix. (**E**) Schematic representation of the cryo-EM data processing pipeline. (**F**) Model fit to map for the whole condensin II pentamer (SMC2 green, SMC4 purple, CAP-D3 lightblue, CAP-G2 pink, CAP-H2 yellow), with zoom in onto the regions with highest resolution. (**G**) Orientation distribution in final set of refined particles. (**H**) Detail of the ABC ATPase domain with the two ADP-BeF3 molecules represented as sticks. (**I**) Detail of the structure and position of the CAP-H2 N-terminal domain sandwiched between the SMC2 neck and the CAP-G2 subunit. (**J**) Upper panel, detail of the CAP-H2 N-terminal domain sandwiched between the SMC2 neck and CAP-G2 subunit in the structure of condensin bound to two ADP-BeF3 molecules. Lower panel, detail of CAP-H2 N-terminal domain rearranged in the condensin II DNA bound conformation.

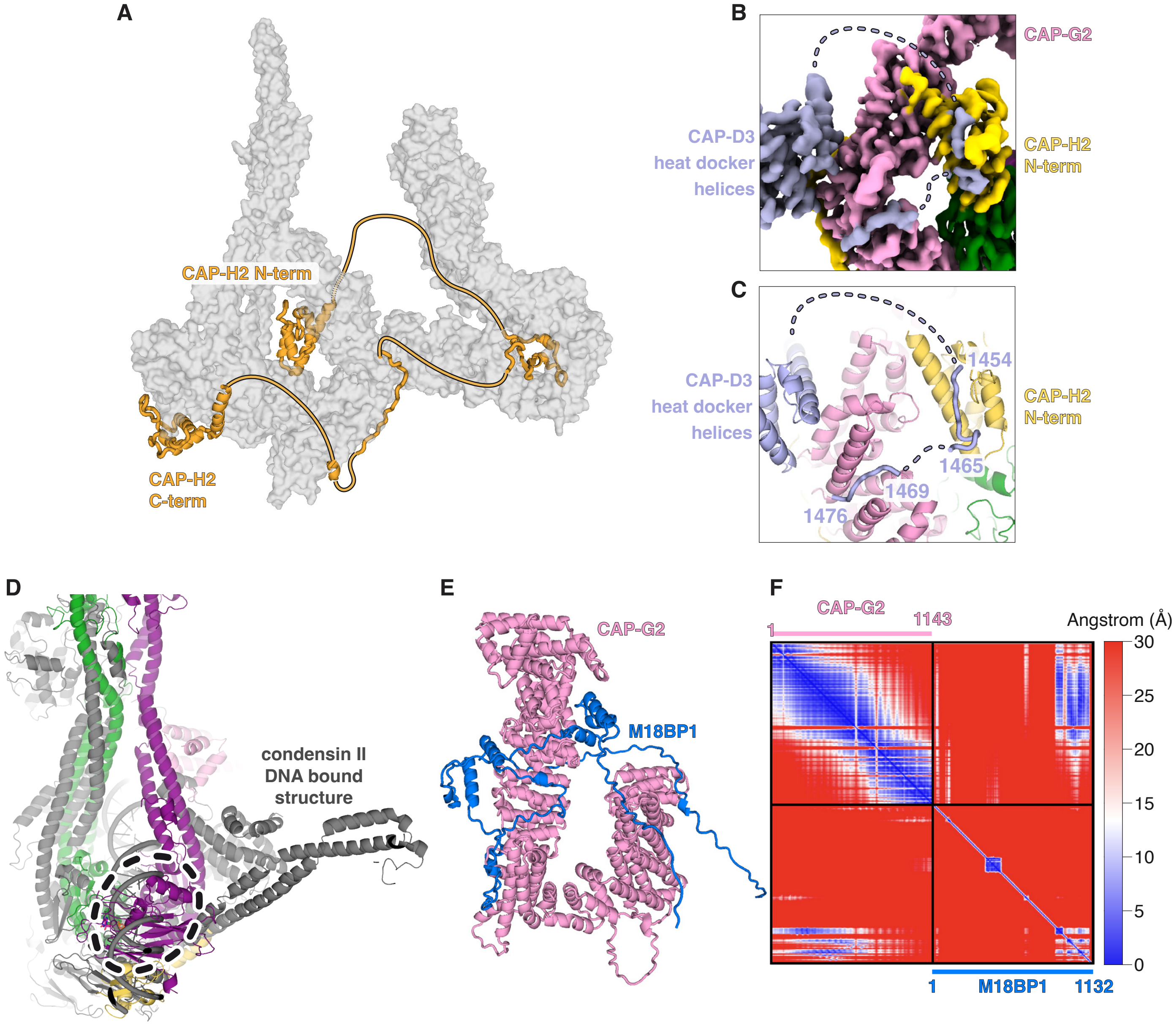


**Fig. S7. Condensin II in the inhibited conformation.** (**A**) Detail of the CAP-H2 interaction sites with condensin II subunits. (**B**) Cryo-EM density map of condensin II boud to two ADP-BeF$_3$ molecules zoomed in the region of interaction of the CAP-D3 C-terminal tail with CAP-G2 and the CAP-H2 N-terminal helices. (**C**) Model of condensin II boud to two ADP-BeF3 molecules, with highlighted residues of CAP-D3 tail that interact with CAP-G2 and the CAP-H2 N-terminal helices. (**D**) Superimposition of condensin II structure in the autoinhibited conformation with condensin II structure bound to DNA colored in grey. Dashed line represents potential clash between the SMC4 subunit of condensin II and the DNA molecule. (**E**) Alphafold3 prediction of the CAP-G2 - M18BP1 interaction, with relative PAE plot in panel (**F**).

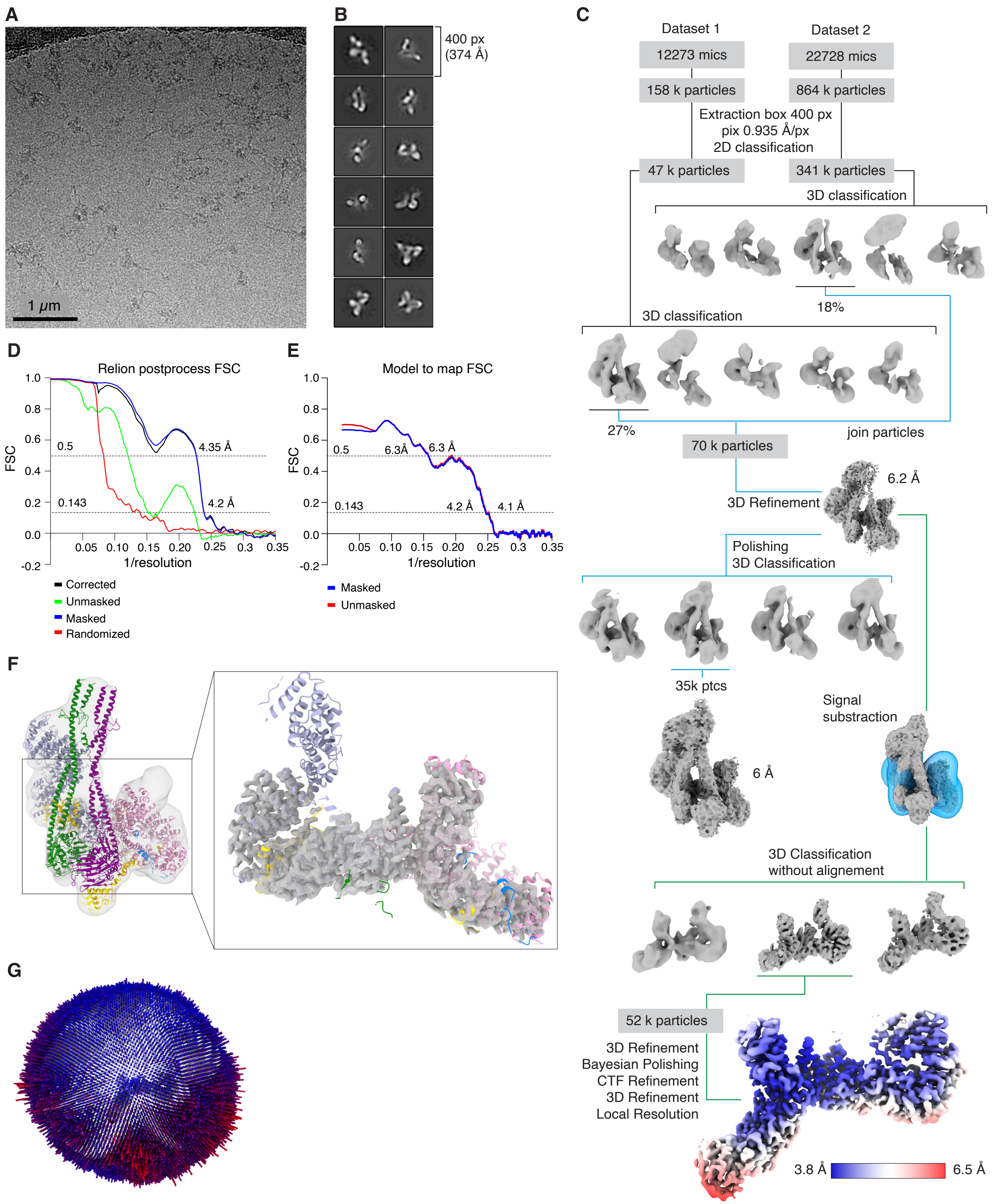


**Fig. S8. CryoEM data analysis of condensin II bound to M18BP1.** (**A**) Representative micrograph. (**B**) 2D class averages from full dataset. (**C**) Schematic representation of the cryo-EM data processing pipeline. (**D**) Fourier Shell Correlation (FSC) curves for raw map obtained with RELION. (**E**) Model to map Fourier Shell Correlation (FSC) curves obtained with phenix. (**F**) Model fit to map for the whole condensin II - M18BP1 complex (SMC2 green, SMC4 purple, CAP-D3 lightblue, CAP-G2 pink, CAP-H2 yellow, M18BP1 blue), with zoom in onto the regions with highest resolution. (**G**) Orientation distribution in final set of refined particles.

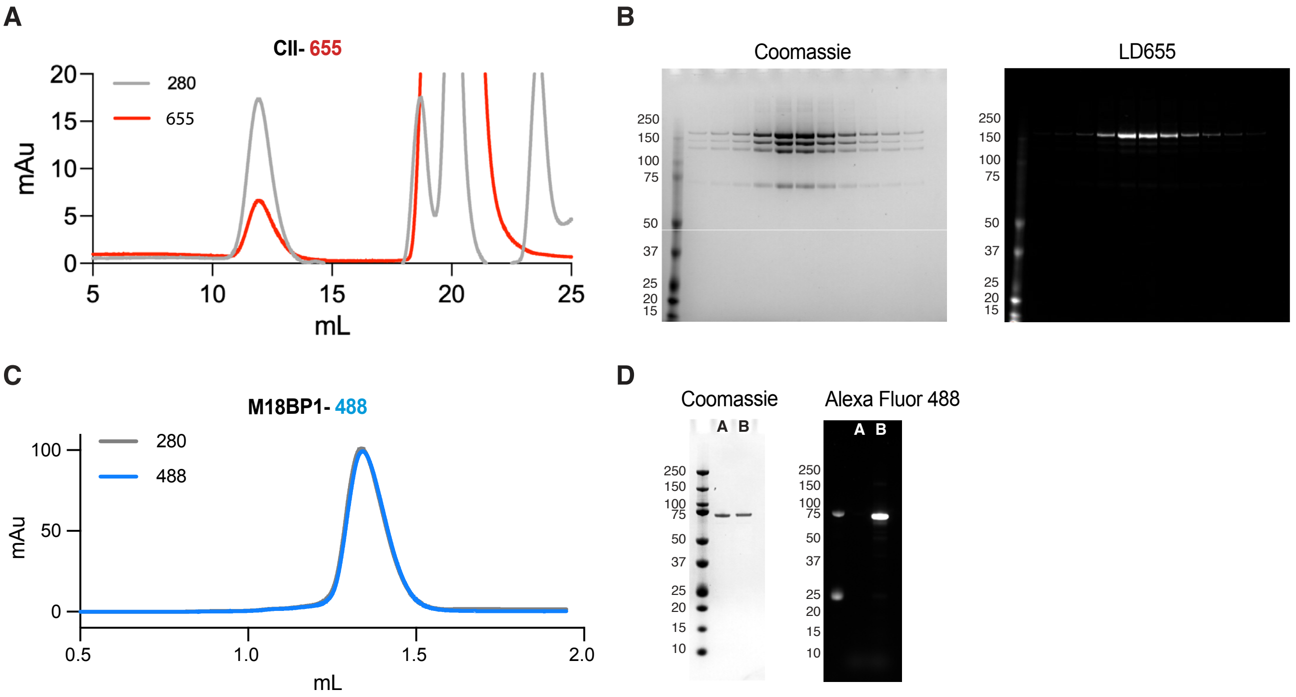


**Fig. S9. Condensin II and M18BP1 labelling.** (**A**) Size exclusion profile (Superose 6 increase, 24 mL volume) of condensin II following labelling reaction with LD655 - CoA and SFP transferase. In grey and red respectively, absorbance at 280 nm and 650 nm. (**B**) SDS page gel of fractions from size exclusion in (a). Left panel, stained with Coomassie blue. Right panel, imaged by exciting at 650 nm. (**C**) Size exclusion profile (Superdex 200 increase, 2.4 mL volume) of M18BP1 following labelling reaction with Alexa Fluor 488 - maleimide. In grey and blue respectively, absorbance at 280 nm and 495 nm. (**D**) SDS page gel of unlabelled (A) and labelled (B) M18BP1. Left panel, stained with Coomassie blue. Right panel, imaged by exciting Alexa Fluor 488.
